# Principles and Operation of Virtual Brain Twins

**DOI:** 10.1101/2024.10.25.620245

**Authors:** Meysam Hashemi, Damien Depannemaecker, Marisa Saggio, Paul Triebkorn, Giovanni Rabuffo, Jan Fousek, Abolfazl Ziaeemehr, Viktor Sip, Anastasios Athanasiadis, Martin Breyton, Marmaduke Woodman, Huifang Wang, Spase Petkoski, Pierpaolo Sorrentino, Viktor Jirsa

## Abstract

Current clinical methods often overlook individual variability by relying on population-wide trials, while mechanism-based trials remain underutilized in neuroscience due to the brain’s complexity. A Virtual Brain Twin (VBT) is a personalized digital replica of an individual’s brain, integrating structural and functional brain data into advanced computational models and inference algorithms. By bridging the gap between molecular mechanisms, whole-brain dynamics, and imaging data, VBTs enhance the understanding of (patho)physiological mechanisms, advancing insights into both healthy and disordered brain function. Central to VBT is the network modeling that couples mesoscopic representation of neuronal activity through white matter connectivity, enabling the simulation of brain dynamics at a network level. This transformative approach provides interpretable predictive capabilities, supporting clinicians in personalizing treatments and optimizing interventions. This manuscript outlines the key components of VBT development, covering the conceptual, mathematical, technical, and clinical aspects. We describe the stages of VBT construction–from anatomical coupling and modeling to simulation and Bayesian inference–and demonstrate their applications in resting-state, healthy aging, epilepsy, and multiple sclerosis. Finally, we discuss potential extensions to other neurological disorders, such as Parkinson’s disease, and explore future applications in consciousness research and brain-computer interfaces, paving the way for advancements in personalized medicine and brain-machine integration.

“*The brain is conceived as a self-organizing system operating close to instabilities where its activities are governed by collective variables, the order parameters, that enslave the individual parts, i*.*e*., *the neurons*.” Professor Hermann Haken (1927-2024).

## 1 Introduction

The Virtual Brain Twin (VBT) may completely change our approach to study and interrogate the human brain and its function. As an example, VBTs might be used for the design of drug discovery and that of brain-computer interfaces. Today, most clinical decisions and approaches are validated via population-based trials. This approach consists of selecting typically thousands of individuals (e.g., for phase III trials), that are randomly assigned to the proposed new treatment or to a different treatment (typically either a sham or the current gold standard). As the vast majority of the clinical trials fail, these are typically followed by retrospective analyses aimed at distinguishing the features of the responders vs. non-responders, which not uncommonly leads the researcher to stumble upon the causal mechanisms by chance [1]. In molecular biology, the realization of the inefficiency of such an approach led to the development of mechanism-based trials, where the efficacy of a drug is tested by selecting participants for multiple diseases that share a molecular pathophysiological mechanism [2]. While this approach was extensively deployed for diseases where the pathophysiological mechanisms were relatively well understood (e.g., cancer), this has not been the case in neuroscience, neurology and psychiatry, where often the pathophysiological mechanisms cannot be expressed entirely in terms of molecular alterations, as brain functions emerge from the coordinated interactions of multiple units [3], at multiple scales and “the total is not the sum of the parts”. In other words, one needs to include a causal model of the mesoscopic activity of brain areas and their interactions to bridge the gap from mechanisms occurring at the microscopic level to symptoms that likely stem from interactions at the intermediate and large-scales [4].

VBTs are digital twins of the full brain network [5, 6], that are constrained by structural brain imaging data and simulate functional sensor data such as electro- and magnetoencephalography (EEG, MEG) and functional Magnetic Resonance Imaging (fMRI). They represent an effective way to encapsulate previous biological knowledge (i.e., functions of channels, effects of neurotransmitters, interactions between excitatory and inhibitory units) in a mathematical representation [7]. Because they are based on fundamental biological principles, these models can extrapolate beyond existing data, and predict the outcomes of interventions that may not have been previously observed [8,9]. This predictive capability is particularly important in the development of new drugs or in tailoring treatments to individual patients, where the model can simulate various scenarios and optimize the treatment plan based on predicted outcomes. Finally, by incorporating key biological or dynamical mechanisms, these models might predict not only whether an intervention will work, but also provide insight about how and why it will (not) work. Hence, large-scale models might constitute a major advancement– conceptual and technical at once–in the study and treatment of brain disorders, in that they can: 1) enhance our understanding and clinical management of neurological and psychiatric conditions; 2) improve and tailor the design of medical instruments for the delivery and monitoring of therapies; and 3) tailor the design of neuromorphic devices to help people with disabilities in everyday life. Furthermore, these models could enable highly personalized treatments for neurological and psychiatric conditions, also in terms of prediction of side effects in individual patients. From a practical standpoint, this capability could extend to optimizing drug therapies [10, 11], tailoring non-invasive brain stimulation techniques, and even designing personalized rehabilitation programs.

Beyond clinical uses, mechanistic brain models have the potential to advance our understanding of fundamental brain functions and cognitive processes [12]. For instance, the dynamics of resting-state networks are crucial for understanding both healthy and pathological brain functions. To diagnose and prognose diseases effectively, it is crucial to investigate the functional organization and emergent dynamics of these networks, and to understand how deviations from normal activity manifest in pathological conditions. They could also play a crucial role in the development of brain-computer interfaces, enhancing communication and control for individuals with motor impairments. Additionally, these models may contribute to the emerging field of neuromorphic computing, where insights into brain organization and processing can inspire new, more efficient computing architectures. The Virtual Epileptic Patient (VEP; [8, 13]), a whole-brain network model tailored to individual epileptic patients, represents a notable use case of VBTs. Simulating the unique brain networks of individual patients, the VEP model significantly improves treatment planning and prognosis for drug-resistant patients who are undergoing surgery [8, 13]. However, these personalized models extend their utility to other neurological and psychiatric disorders. For this translation to occur, mechanistic and phenomenological models are built upon a detailed understanding of the physiological processes and structures involved. For instance, in the context of healthy aging, these models track the natural changes that occur in brain networks over time [14]. In Alzheimer’s disease, brain models have been built to help researchers understand the disease’s progression and its impact on the brain networks [15]. For diseases like multiple sclerosis and Parkinson’s disease, the models provide valuable insights into how these conditions affect brain connectivity and function [16, 17]. In the realm of psychiatric disorders, great promise is held by the use of large-scale models to link the effect of neurotransmitters on large-scale brain dynamics [18].

In this manuscript, we will guide the reader through all the main building blocks– conceptual, mathematical, technical, and clinical– leading to the construction of a VBT. In section 2, we will lay out the main ideas on which a VBT is built, focusing on the digital twin cycle, its components, and providing a general overview. In section 3, all the building blocks of the VBT will be described in detail, covering commonly used approaches to neural mass modeling as a key component, along with alternatives such as data-driven techniques. Furthermore, emphasis will be put on the use of anatomical coupling based on the brain connectome, as a way to achieve (low- and high-resolution) personalization of large-scale models, along with the associated mapping from sources to sensors. Then, in section 4, we will provide an overview of the stages involved in operating VBTs, from the preprocessing of structural data, constructing brain atlases and coregistration in the subject-specific brain space, to simulation and inference, that is how to confront VBTs with functional data to estimate quantities of interests (i.e., the control parameters for personalization). Finally, in section 5, we will demonstrate the use cases of VBTs (including resting state, healthy aging, multiple sclerosis, and epilepsy), with an emphasis on their contributions to advancing clinical practice.

## 2 Concepts of Virtual Brain Twin

The Virtual Brain Twin (VBT) finds its roots in nonlinear dynamics, self-organization and synergetics [19, 20]. First efforts of building full-brain network models date back to the early 70s, when Paul Nunez discussed brain waves and global wave properties of EEG using dispersion relations, linking spatial wave lengths to temporal frequencies as observed in scalp recordings on the *cm* scale [21–23]. He linked the dispersion relation to the brain’s network connectivity, specifically the exponential decay of fiber length over distance. Nunez’ considerations were entirely based on rodent data on the 1 *cm* scale, which he then scaled up to the human on the 10 *cm* scale. Nunez used this theoretical framework to quantitatively explain several features of rhythms observed in EEG. Instead of modeling multiple populations, as in the Wilson-Cowan [24] and Amari [25] field equations, the Nunez brain wave equation effectively models a single population with both excitatory and inhibitory synapses [26]. Notably, Nunez was amongst the first to integrate time delays via signal propagation into his brain wave equation, which were necessary, considering the macroscopic scale of *cm* [27]. The approach was fully linear, although the necessity of nonlinear pattern forming mechanisms was acknowledged. In 1996, Jirsa & Haken [28, 29] provided such a nonlinear formulation and derived a brain wave equation, which considered the split of the brain connectivity into the intracortical short-range fibers and the corticocortical long-range fiber system, which later became known as the connectome. Both fiber systems included time delays via signal propagation, albeit with different speeds. The Jirsa-Haken equation combines the linear spike-to-wave conversion at the synapses with the nonlinear wave-to-spike conversion at the cell bodies (see Walter Freeman’s work on neural masses [30–32]) and presents them in a closed form. Its initial formulation followed Nunez’s approximation, in which the long-distance connectivity was modeled using an exponentially decaying integral kernel. In collaboration with Scott Kelso [33,34], however, it became rapidly evident that many basic phenomena known from large-scale brain dynamics cannot be explained, not even as an approximation, by a spatially invariant large-scale connectivity (Figure 1**A**).

**Figure 1:**
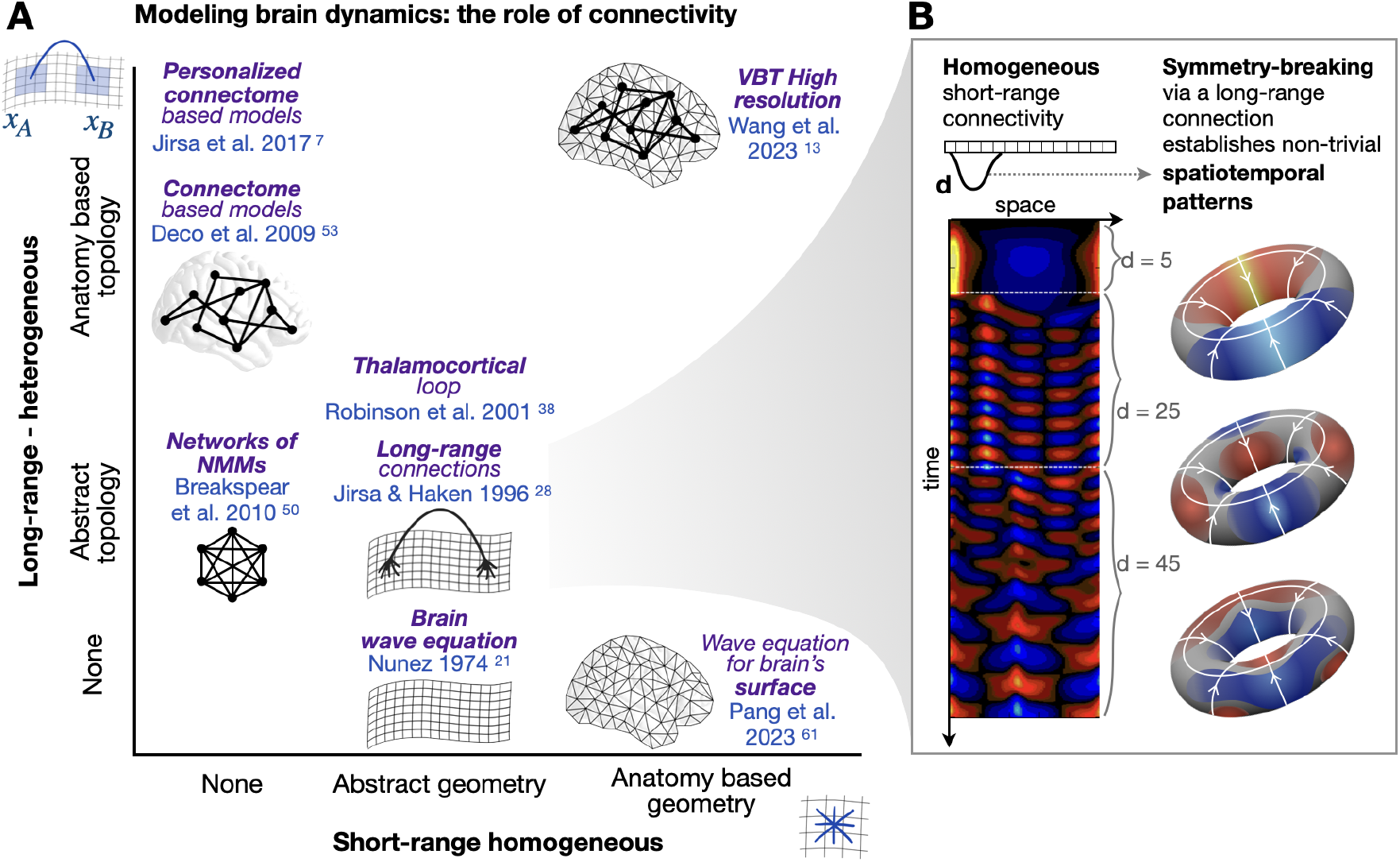
The role of homogeneous and heterogeneous connectivity in the development of models for brain activity. (**A**) In the course of developing brain models, the focus has shifted from homogeneous connections between neighboring nodes on the cortical surface to the long-range connections among distant nodes, as described in the connectome. Technological advances have now enabled the construction of personalized brain models, incorporating both long- and short-range connectivities. These approaches have converged in high-resolution VBTs. Here, we have cited some exemplary references to illustrate key concepts. (**B**) Symmetry breaking of connectivity through long-range connections plays a key role in shaping non-trivial spatiotemporal dynamics. This is demonstrated in a simplified model with one dimension and a single long-range connection of length *d*. During the simulation, *d* is varied, and the system settles into new patterns. The structure of these periodic patterns is illustrated on toroidal manifolds.

To become more concrete, let *ψ*(*x, t*) be the neural vector field of dimension *N* capturing the population activity at time point *t* and position *x*. The dynamics of the neural field can then be described by the following integro-differential equation [28, 35]

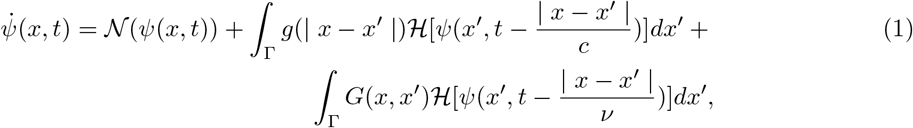

where 𝒩 (*ψ*(*x, t*)) is the local population model of dimension *N*, with *N* typically ranging from 1 to 12, depending on the neural mass model used. The spatial domain of the neural field is denoted by Γ, which can be one, two, or three dimensional. In the simplest one-dimensional case, *x* ∈ Γ = [0, *L*] and *L* is the spatial length of the neural field. The homogeneous (short-range) connectivity function, *g*(| *x* − *x*^*′*^ |), is translationally invariant. If the connectivity function does not exhibit this property, it is referred to as heterogeneous, denoted as *G*(*x, x*^*′*^)≠ *G*(| *x* − *x*^*′*^ |). Heterogeneous fibers are typically long-range and are myelinated, establishing the white matter of the brain. The parameters *c* and ν represent the propagation velocities through the homogeneous and heterogeneous connections, respectively.

To illustrate some of the properties of networks with time-delayed mixed fiber systems, assume 2*σ* the local neural mass dynamics to be linear, 𝒩 (*ψ*(*x, t*)) = −*ϵψ*(*x, t*), and the cortical surface to be one-dimensional with connectivity 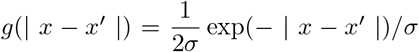 (i.e., finite propagation speed, with translationally invariant connectivity). Using the method of Green’s functions [36] and applying the inverse Fourier transform, we can rewrite the above integral equations as a nonlinear partial differential equation for the field [28, 29]

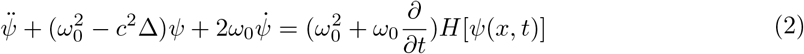

where *ω*_0_ = *c/σ* > 0 represents a characteristic frequency, and Δ denotes the Laplacian. In the absence of input, the left hand-side is a damped wave equation with oscillatory properties. The spatially uniform pattern tends to remain stable, if the slope of the sigmoid function *H*[·] on the right-hand-side is sufficiently small, though transient wave propagation may occur following small discrete perturbations [37]. If the slope exceeds a certain threshold, the spatially uniform state becomes destabilized and wave propagation may occur. Based on Robinson’s approach [38, 39], analytic and numerical predictions for EEG spectra in terms of physiological parameters have shown agreement with empirical EEG data, considering a subcortical feedback loop that may underlie the production of the alpha rhythm [40], such as feedback through the thalamocortical projection system [41–44].

In absence of heterogeneous fibers, at least in an approximate manner, any activity generated at a location *x*_*A*_ would have to propagate through all intermediate zones to generate an activity at a distant location *x*_*B*_. In the other extreme, all brain tissue is parcellated into regions with no homogeneous connectivity. Each region contracts into a point-like neural mass, compressing all information to a maximum. The outcome is a network of nodes, which are connected by the spatially variant long-range fiber system where, *G*(*x, x*^*′*^)≠ *G*(*x* − *x*^*′*^). The importance of the symmetry breaking cannot be overemphasized, especially for large-scale brain networks [45–48]. It recognizes that, first, the topology of the network architecture is fundamental to the spatiotemporal pattern formation (Figure 1**B**), and second, the connectivity has a space-time structure, as each link of the network is characterized by a weight and a time delay relating two brain regions [49]. Breakspear et al. [50] pointed out that while global (all-to-all) coupling amongst system oscillators (e.g., Kuramoto model [51] or generic Hopf [52]) may be a reasonable approximation for small, densely connected neural networks, it is certainly not true for large populations distributed across the cortical sheet. Instead, coupling should be spatially embedded, incorporating time delays between distant subsystems and reducing coupling strength with distance. Deco and colleagues [53, 54] have also emphasized that resting-state networks emerge from a dynamic framework of noise, anatomical connectivity, and time delays. The importance of the topological component of connectivity is widely recognized nowadays [55, 56], evidenced by the many publications using graph theory and its toolboxes in neuroscience applications [3, 57–60]. A recent work by Pang et al. [61] further highlights this growing trend by deriving geometric eigenmodes (cortical shape) that account for the effects of local connectivity that is exponentially decaying over the geodesic distance of the cortex, i.e., the exponential distance rule [62–65]. The recognition of the consequences of the temporal component of connectivity is more challenging, but slowly becoming recognized. Petkoski and colleagues, for example, have demonstrated that the space-time structure of the connectivity spans a resonance body, in which brain network activity shows spectral preferences, necessitating the reformulation of traditional graph theory to account for time delays due to finite transmission speeds, and to derive a normalization of the connectome [66, 67]. When applied to the Human Connectome Project (HCP) database, this approach explains the emergence of frequency-specific networks, including the visual and default mode networks. These findings are robust across human subjects (*N* = 100) and are a fundamental network property. Although theoretically intriguing, the actual importance of the connectome’s space-time structure in real-world applications still needs to be demonstrated. That said, the predictive value of the connectome in personalized medicine is well demonstrated. Jirsa et al. [8] have introduced personalized connectome based models to predict the outcome of surgical interventions, and assist in the clinical decision-making process through virtual surgeries [5]. Wang et al. [13] have presented a digital workflow with both low- and high-spatial resolution for mapping individual epileptogenic zone networks based on patient MRI and SEEG recordings.

At this point, it is worthwhile to pause and consider the full brain network from a theoretical perspective, returning to our initial quote by Hermann Haken, “The brain is conceived as a self-organizing system operating close to instabilities where its activities are governed by collective variables, the order parameters, that enslave the individual parts, i.e., the neurons” [68]. Indeed, brain network activity evolves spatiotemporally within a framework, in which the nodes of the self-organizing system experience each a different pattern of interactions, that is the set of all incoming signals communicated via short- and long-range fibers, and the set of all outgoing signals. In- and out-going interactions are not identical, as the connectivity matrix is not symmetric. Mathematically, such interaction pattern translates into spatially variant interaction kernels, which are unknown in traditional physics based on translationally invariant forces. Other networks such as airline traffic networks maybe cited to support a similar network topology comprising short- and long-range connections with the organization of network hubs, but they do not support the oscillatory wave nature of network activity as the brain does (they communicate in a particle mode, flying airplanes from one node to another). The brain’s communication network thus appears to be a unique self-organizing system in this regard. Following Haken’s thinking, when operating close to instabilities, in other words close to a transition from one state to another mediated by a bifurcation, low-dimensional dynamics arises in a self-organized manner. The mathematical basis is the local center manifold theorem, which explains time scale separation in the neighborhood of instabilities and provides a mechanism via adiabatic elimination to enable dimension reduction. Haken called this process enslaving, in which sets of state variables operating close to the instability show a slow dynamics, whereas the fast variables in the residual high-dimensional space are guided towards a manifold following a flow determined by the slow variables, the so-called order parameters. The time scale separation into fast and slow variables has been at the core of this mechanism underlying pattern formation. Over the past years, manifolds have found conceptual entry in the neuroscience literature, where neuronal dynamics have been visualized on manifolds using various dimensionality reduction techniques [69–74]. It has been argued that the role of manifolds in neurosciences goes beyond mere visualization, and that dynamics on manifolds carry computational and functional meaning. It needs to be established how the dimensionality reduction occurs in the brain, with candidate mechanisms including low-rank connectivity [75], averaging and decorrelation [45]. Jirsa & Sheheitli [45] demonstrated the emergence of structured flows on manifolds (SFM) through symmetry breaking in the low-dimensional space [76–78]. Dynamical systems with symmetries are called equivariant systems. Specifically for large-scale networks, it is intriguing to explore how symmetry breaking relates to the connectome. As the connectome changes during the lifespan, from neurodevelopment to neurodegeneration, and in brain disease, the formation of structured flows constrained to manifolds will be affected by these processes [13, 14, 78]. The theoretical framework of SFMs provides a formal language in terms of dynamics to describe the effects of changes in connectivity and network nodes. In other words, the SFM is the mathematical object pertaining to the dynamics of the self-organizing process in the brain network, sufficiently general to capture multistable network states, limit cycles, travelling waves or any low-dimensional attractors. During inference, the SFM is the mathematical object sampled by the inference process. The SFM thus serves, on the one hand, to describe mechanistically spatiotemporal pattern formation processes, and, on the other hand, as a target for causal inference when investigating empirical brain imaging data.

The Virtual Brain Twin (VBT) embraces both of these facets of use of SFMs, the mechanistic forward simulation and pattern generation of the large-scale brain network, and the inference using the brain network model for causal hypothesis creation. The dual use of brain modeling (simulation and inference, forward and backward) has been compulsory for the successful clinical translation and personalized medicine, establishing the workflow associated with the use of VBT. In closure of this conceptual section, we summarize the key ingredients of the VBT: it is the digital representation of an individual’s brain as a full-brain network model, which integrates the same individual’s structural and functional brain imaging data in the model building (hence “Twin”). The process of creating a VBT comprises three stages (see Figure 2): first, structural brain imaging data (MRI, DTI, eventually CT) are co-registered in the same space. MRI data are parcellated into nodes, which, depending on the desired resolution, may vary from 10 *cm*^2^ and hundreds of nodes, to 1 *mm*^2^ and hundred thousands of vertices. In the former case, a brain region typically equals a node, whereas in the latter case a brain region spans hundreds of vertices. Tractography is performed from the DTI data to create the individual’s connectome. The nodes are connected through the connectome to a network, which is spanned in physical space. This step is critical, as it allows to position correctly the brain imaging sensors in the same space. Depending on the brain imaging modality, sensors are EEG contacts, MEG SQUIDs, and voxels in fMRI. The source-to-sensor mapping is computed from the relevant physics equations. Second, each node is equipped with a neural mass model, which is a reduced mathematical representation of the dynamical equations prescribing the behavior of large populations of neurons, typically at least several hundred thousands. These large numbers are necessary to justify the use of statistical methods, leading to the dimension reduction. Neural masses still maintain the multiscale nature of the processes ongoing in a neuronal population such as fast neuronal discharges and the slower chemical processes such as extracellular potassium changes. The connectome provides the connectivity matrix linking the activity across nodes through white-matter tracsts. At high spatial resolution, the vertices in a brain region integrate the connectivity within the cortical areas in addition to the highly heterogeneous connectivity between the brain regions, the so-called connectome. Third, using functional brain imaging data, the application of Bayesian inference methods provides an estimation of relevant model parameters. Typically, not all parameters are inferred. Most of them are set into operating ranges from the literature based on physiological estimates or mathematical calculations. The only inferred parameters are those that are relevant to the intended use of the VBT, typically linked to some pathophysiological cause in clinical applications, for instance the epileptogenic zone in epilepsy or lesions in multiple sclerosis. This approach supports the convergence of the inference process.

**Figure 2:**
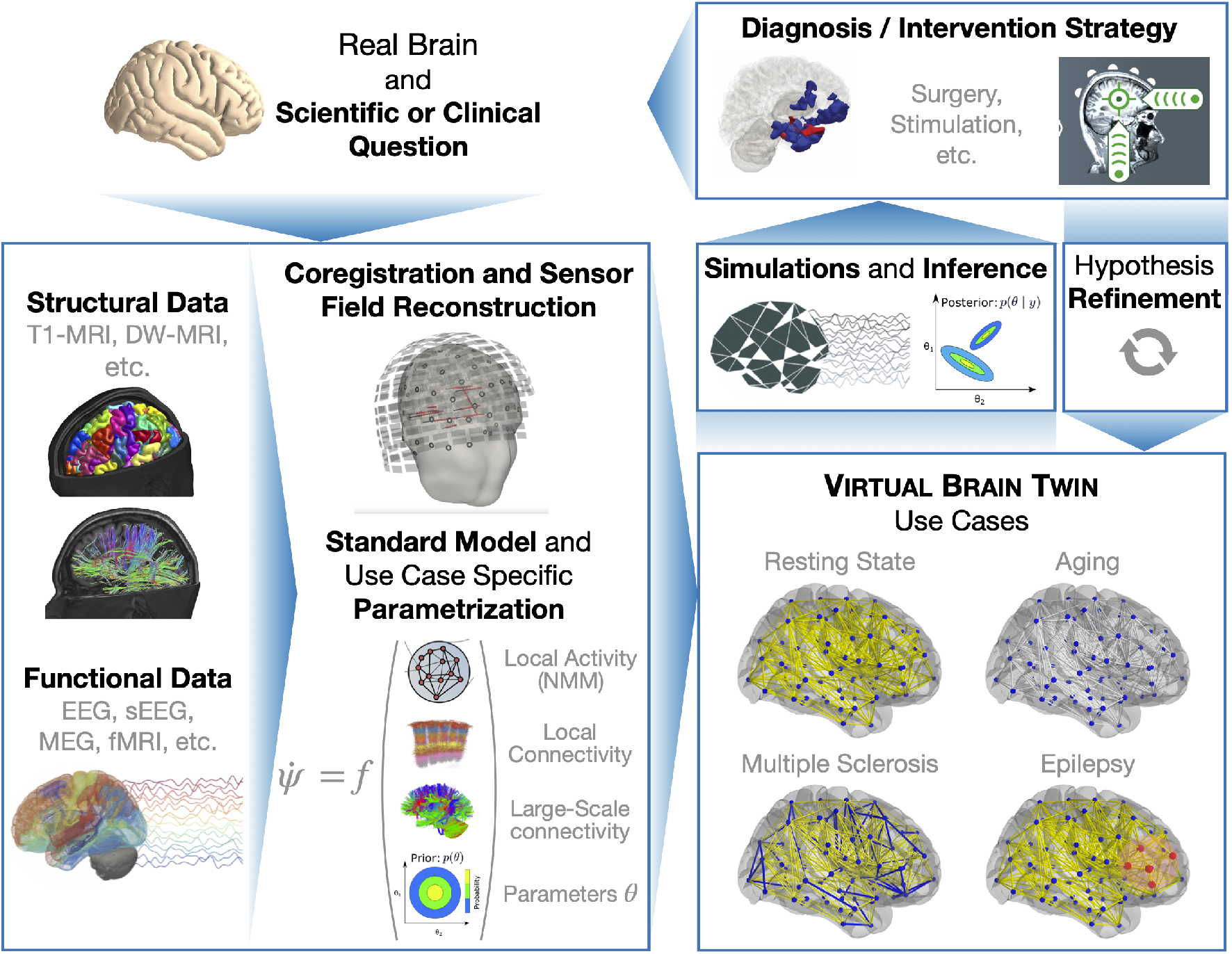
Workflow to create a Virtual Brain Twin (VBT): Based on scientific questions or clinical applications, appropriate structural and functional clinical data enable the parametrization of a standard model. Along with source reconstruction, this allows for the creation of a VBT in which hypotheses can be tested through simulations and inferences, aiding in diagnosis and interventions.

## 3 Building the Virtual Brain Twin: nodes and connectome

In the following sections, we will describe the process of building the VBT, covering the construction of nodes and the connectome. We will provide an overview of various neural mass modeling approaches, including biophysical, phenomenological, and data-driven models. Additionally, we will formulate connectome-based models at low- and high-resolutions and outline the methodology for mapping from source level to sensor measurements.

### 3.1 Neural Mass Models as network nodes

Neural mass models (NMMs) represent the activity of large populations of strongly interconnected neurons, corresponding to the mesoscopic scale, which lies between the microscopic cellular level and macroscopic whole-brain models [79]. These models capture the population dynamics by describing the evolution of a few collective mesoscopic variables through the use of differential equations. This formidable dimensionality reduction is the first key feature of NMMs, making it feasible to simulate whole-brain activity and develop personalized large-scale models for clinical applications [7, 80].

Moving from a fine-grained to a more coarse representation entails a series of assumptions specific to the approach that is used (see Table 1) and results in tremendous information compression [79, 81]. However, this approach has the intrinsic advantage of focusing attention on the emergent properties of brain activity that are characteristic of the mesoscopic scale of description. The second key advantage of using NMMs is, thus, that of improving our scientific understanding at the scale for which we have direct access through several measurement techniques in human subjects, such as EEG, MEG or fMRI. Empirical recordings of brain activity at the whole brain level are only possible with very limited spatial (or temporal) resolution. High spatial resolution can be achieved using fMRI, which is capable of reaching a ∼ *mm*^3^ resolution [82, 83]. Even in this best-case scenario, it is estimated that a single fMRI voxel contains ∼ 500*k* interconnected neurons [84], which form complex interactions on their own, making it intractable to infer the causal links between neuronal organization and whole-brain activity. Consequently, the coarse lens through which we observe neuronal activity at the whole-brain level supports the use of NMMs.

**Table 1:** Examples of commonly used Neural Mass Models (NMMs), in the context of VBT.

| Model | Derivation | Timescales | Repertoire | Number and Interpretation of Variables |
| --- | --- | --- | --- | --- |
| **Wilson-Cowan, 1972 [24]** | A2 - first mesoscopic description of interacting excitatory and inhibitory populations | fast | up/down, limit cycle | #2; firing rates of two populations (Excitatory and Inhibitory) |
| **Jansen-Rit, 1995 [89]** | A2 - extends Wilson-Cowan model to three populations | fast | up/down, coexisting limit cycles (related to high alpha and low beta EEG oscillations) | #6; membrane potentials and firing rates of pyramidal neurons, excitatory, and inhibitory interneurons |
| **Wong-Wang, 2006 [90]** | A3 - reduction from biophysical model of interaction of excitatory and inhibitory populations | fast | multi-stability | #2; synaptic gating variable of the excitatory and inhibitory populations |
| **Stefanescu-Jirsa 2/3D, 2008 [91]** | A1 - dispersion of parameters causes clustering of neurons (FitzHugh-Nagumo / Hindmarsh-Rose) [92] | fast, slow/fast | up/down, limit cycle/bursting | #2; or #3; state variables representing excitatory and inhibitory interactions |
| **Destexhe et al., 2009-2024 [93–96]** | A1 - based on transfer function obtained from several neuron models or recordings | fast, slow/fast (with adaptation) | up/down, additional stable fixed point | #3 (#6; with 2nd order); fast-spiking inhibitory and regular-spiking excitatory firing rates + adaptation (+ 2nd order) |
| **Epileptor [97]** | A4 - slow variable mechanisms to start and stop seizures through specific bifurcations to produce precital spikes and spike-and-wave complexes through an excitable system | fast, intermediate, slow | up/down, limit cycle, bursting | #6 field potential as a combination of phenomenological variables acting on different timescales |
| **Montbrió et al., 2015 [98]** | A1 - starts from a network of QIF neurons, exact assuming the Lorentzian ansatz | fast | up/down | #2; mean-membrane and firing rate of an excitatory population |
| **Depannemaecker et al., 2024 [99]** | A1 - extends the aQIF-MF model (Chen & Campbell [100]) to include dopamine dynamics | fast, slow, ultra-slow | up/down, bursting | #8; firing rate, mean membrane potential, adaptation, inhibitory and excitatory synaptic activations, dopamine receptor activation, local dopamine concentration |
| **Generic 2D [101]** | A3-A4 - extends the dynamics of single neurons (FitzHugh-Nagumo Model [102, 103], Morris-Lecar [104]) | slow/ fast | up/down, limit cycle | #2; membrane potential, and a recovery or damping variable |

We take the pragmatic approach to modeling phenomena at a macroscopic or mesoscopic scales, particularly when the microscopic details are either not fully understood or are deemed irrelevant in the context of the empirical data. One key advantage of this strategy is its ability to generalize across systems where different microscopic configurations can lead to the same emergent global dynamics. This is called degeneracy [85], which is the geometrical counterpart of non-identifiability. In other words, while the microscopic realizations may differ, they do not necessarily affect the system’s overarching behavior, enabling the development of models without the need to precisely capture every fine detail. This focus on the mesoscopic scale also aligns well with the practical considerations of clinical applications. In translational contexts, the mesoscopic scale is the most accessible and measurable, making it the natural target for data collection and analysis. By prioritizing this scale, models can be directly applied to clinical settings, offering actionable insights while bypassing the complexity and potential noise of microscopic-level details.

There are several routes to designing NMMs, each providing unique insights into the modeling of brain activity (Figure 3**A**). When specific biological assumptions and constraints are incorporated, ensuring that the model’s variables maintain an explicit link to the underlying physiology, we refer to these as biophysically inspired or mechanistic models [86]. When the focus is rather on directly reproducing the phenomenon of interest and making hypotheses about the underlying dynamics, we deal with phenomenological models of abstract variables and parameters. These dynamics can also be inferred through data-driven machine learning approaches (see subsubsection 3.1.4). Indeed, there exists a continuum of different models, ranging from the most mechanistic and detailed model to the most reduced and phenomenological one. The different components of the model can encode different levels of biophysical or phenomenological description, providing more detail on the mechanisms of interest for a given study or application. This variety in modeling approaches allows to choose the appropriate level of abstraction based on specific needs [87]. In addition, phenomenological models can identify dynamical structures that can be related and mapped onto the behavior of more detailed biophysical models [87,88]. In this way, they build the bridge between biological mechanisms and emergent dynamics (Figure 3**B**). In the following, we will describe these approaches in more detail.

**Figure 3:**
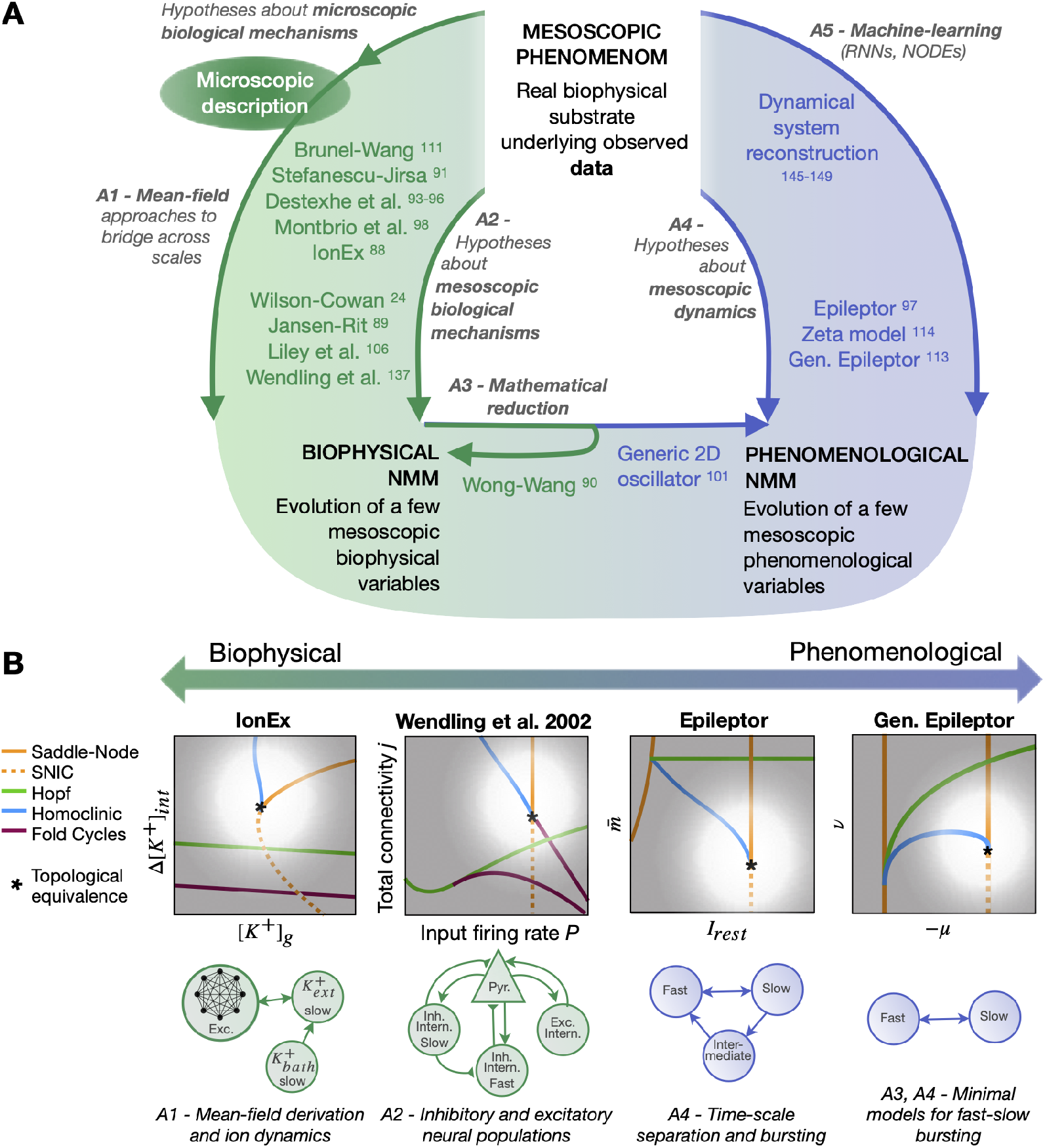
Neural Mass Models (NMMs). (**A**) Main approaches to build NMMs, with specific models applying them at varying degrees and lying on the continuum between biological and phenomenologically inspired. For biophysical models, hypotheses can be cast also at the microscopic level and be followed by the application of mean-field methods to derive the mesoscopic model (A1), or the hypotheses can be advanced directly at the mesoscopic scale (A2). Biophysical NMMs can also be obtained through the mathematical reduction of more complex models (A3). The latter approach can be used to obtain a phenomenological model reproducing key dynamical features of a biophysical one (A3). Another route to phenomenological models is to start directly from data and implement hypotheses about the mesoscopic dynamics (A4) or by applying machine-learning approaches (A5). This leads to models with different characteristics (see Table 1) tailored to specific applications. (**B**) Despite this heterogeneity, models can share some core dynamical mechanisms, allowing for a bridge among them. Here we show examples from seizure modeling. A specific structure in their bifurcation diagram (top row, white region surrounding the asterisk), enables seizures in all these models. However, the approaches to modeling are quite diverse, as is the interpretation of variables and parameters.

#### 3.1.1 Biophysical inspired models

The mechanisms of biophysically inspired models can be directly motivated at the mesoscopic scale, as in the seminal Wilson-Cowan model of two interacting excitatory and inhibitory neural populations [24], and its extensions [89, 105, 106], inspiring transfer function-based mean-field approaches [43, 93, 95, 96]. The model’s equations can alternatively be derived by applying mean-field theory to networks of coupled neurons [26]. Under a series of assumptions, this approach yields the behavior of the mean of the population activity and, potentially, of the higher momenta [79]. A special class of these models, referred to as next-generation NMMs [107,108], relies on the assumption of all-to-all connected neurons with heterogeneity following a Lorentzian distribution. This approach provides an exact derivation for infinitely large networks (in thermodynamics limit, i.e., *N* → ∞) [98, 100, 109], and accounts for the degree of within population synchrony. The original approach developed by Montbrió et al. [98] for population of quadratic integrate-and-fire (QIF) neurons, and has been extended to account for physiological factors such as adaptation [100], or ionic exchange [88]. To exemplify some key features of NMMs, we briefly describe the first of these next-generation NMMs, given by [98], which is a minimal representation upon which the other derivations add extra complexity and biological realism. The model reads:

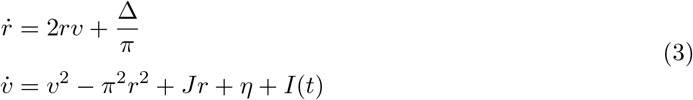

where *r*, and *v* represent the mean firing rate and the mean neuronal membrane potential of the population, respectively. The parameter *J* denotes the synaptic weight, *η* is the average excitability, and Δ indicates the spread of the neuronal excitability distribution in the neural population.

The NNM of Montbrió et al. (given by Equation 3) imposes a flow in the low-dimensional space spanned by the system’s variables (i.e., how the system’s dynamics evolve in state space) (Figure 4, top). For certain rameter settings, the model demonstrates bistability, characterized by the coexistence of a lower firing rate regime (down-state) and a higher firing rate regime (up-state). This feature, the coexistence of two attractors in state space, is common to several NMMs [24,89–91,93,97,106,110–115], and is crucial for capturing key characteristics in empirical data, such as functional connectivity dynamics and neuronal cascades [116,117]. The up-state often facilitates oscillatory activity, which can be either damped, as seen in the next-generation NMMs [98], where the up-state is a spiral fixed point, or sustained when it forms a limit cycle [24, 91, 97, 118]. Some models even allow for the coexistence of limit cycles in addition to the down-state, accounting for the presence of different frequency bands in the EEG [89, 106] (Figure 4, bottom). Transitions between coexisting attractors can occur due to noise or inputs to the NMM. Changes in the parameter values can cause a modification in the number or types of attractors, leading to a qualitative change in the system’s dynamics. In such cases, the system is said to undergo a bifurcation, which introduces another mechanism for transitions. The two mechanisms, noise induced transitions and parameter changes causing bifurcations, are shown in Figure 4 (top and bottom, respectively).

**Figure 4:**
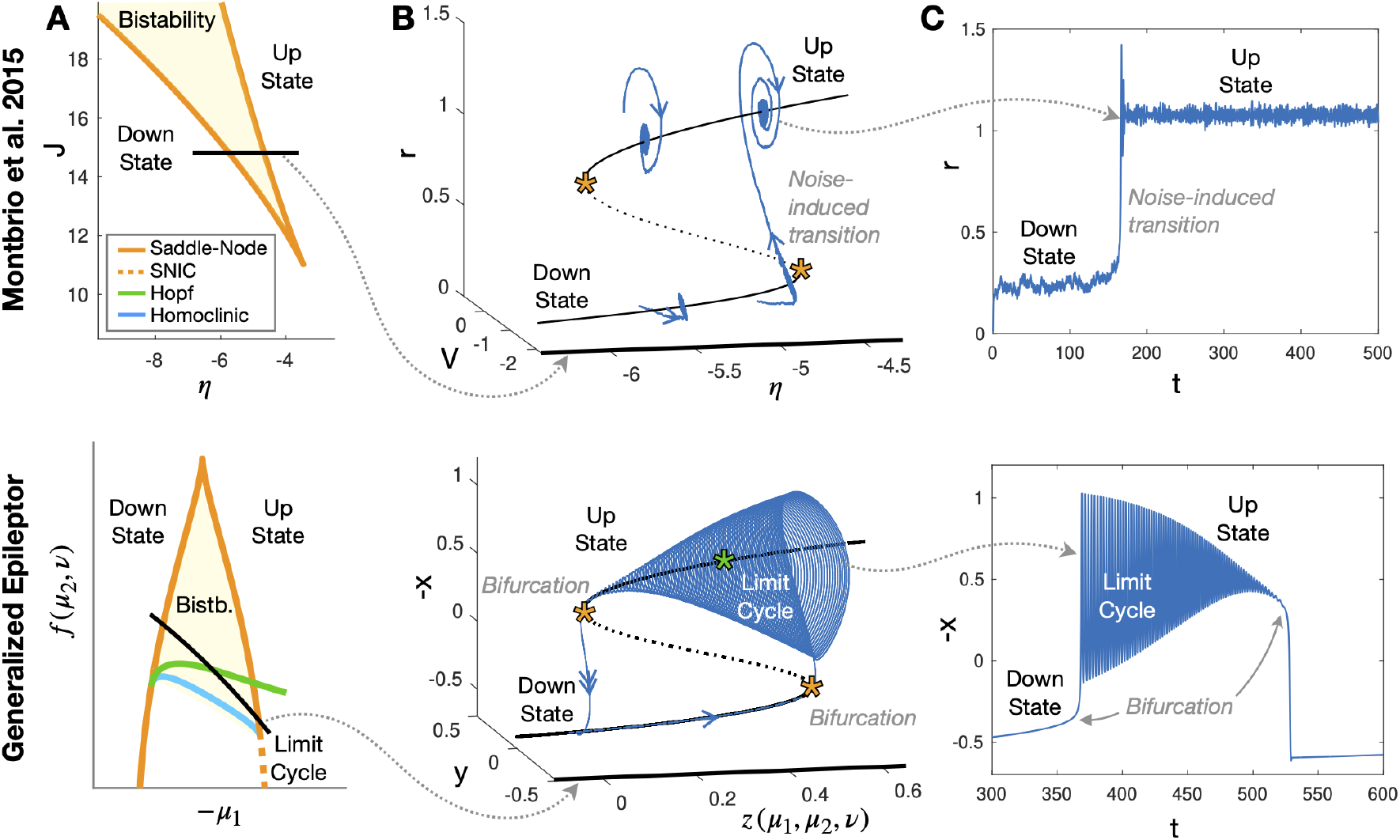
Examples of time series and bifurcation diagrams in NMMs. (**A**) Time series in NMMs showing transitions between states. In the model of Montbrió et al. [98] (top), transitions are induced by noise when the system is near a saddle-node bifurcation. In the Epileptor model and its generalized version [110] (bottom), transitions are driven by slow variable dynamics, which push the system across bifurcations (saddle-node in the figure, although additional bifurcations can be accommodated in these models). **(B)** The manifold of equilibria for the two models (solid black line for stable fixed point, dashed for unstable, and dotted for saddle) when varying one parameter. The time series are overlapped, including examples of orbits in the Montbrió model that, being too far from the saddle-node, do not exhibit transitions. For this representation, we set Δ = 1 and *J* = 15. **(C)** Bifurcation diagrams. In both cases, saddle-node curves organize the bistability between fixed points. The Epileptor, however, exhibits additional bifurcations that produce a limit cycle in the up-state allowing for sustained oscillations.

**Figure 5:**
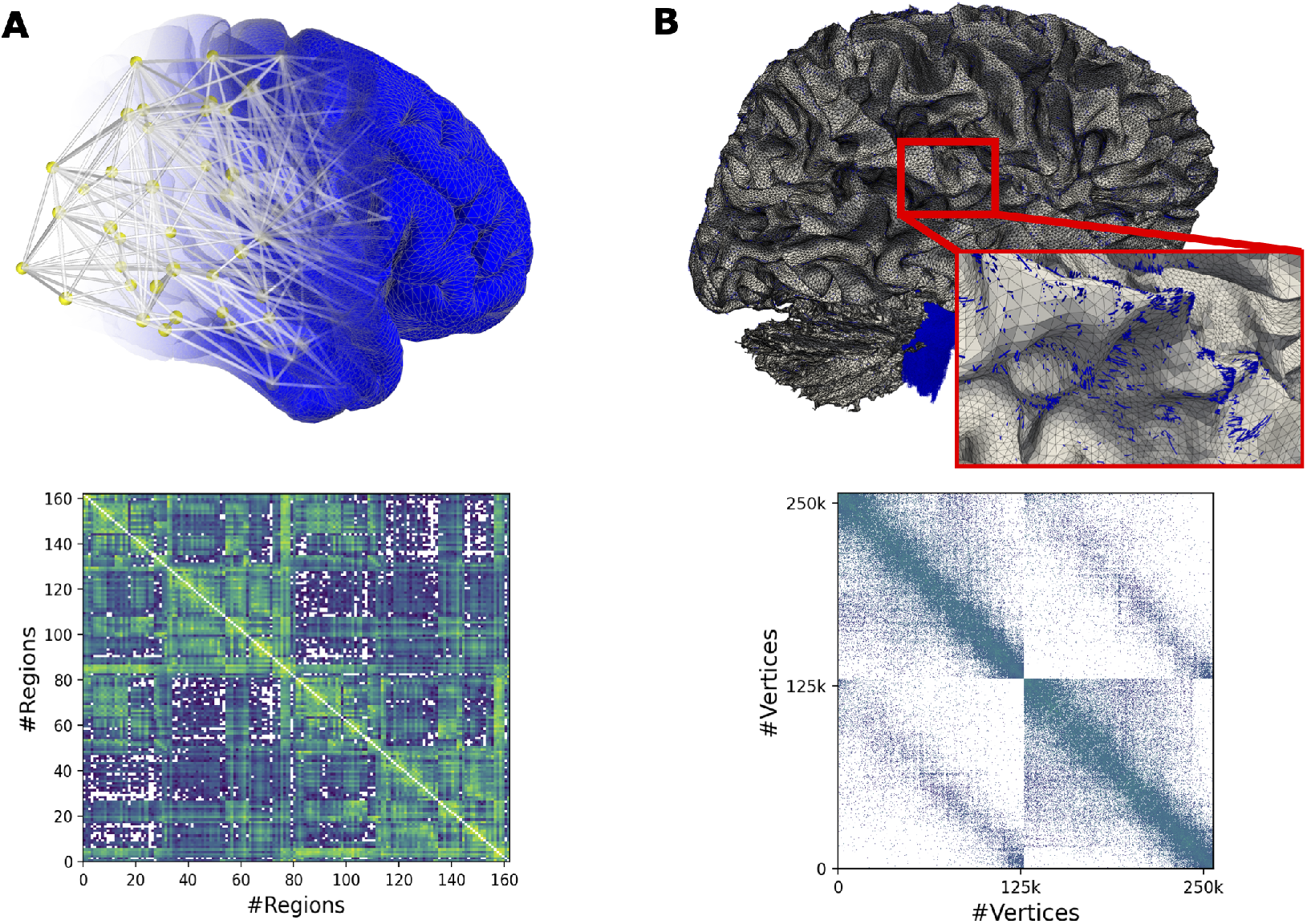
The low- and high-resolution simulations in VBTs. (**A**) The network with a neural mass model at low resolution, showing the 3D brain network (top) and connectivity between 162 regions (bottom). (**B**) The network as a neural field model at high resolution, showing cortical surface with streamline intersections (top) and derived connectivity between 260k vertices (bottom).

#### 3.1.2 Phenomenological models

The complete set of attractors and bifurcations, that is the bifurcation diagram, provides a powerful description of the dynamical repertoire of a model. Models derived from different approaches often share core properties of the bifurcation diagrams, such as the bistability between down-state and up-state described earlier, identifying these as key features of NMMs. When dynamical systems share these properties, they display the same variation of phase flow in the state space, which allows to find a homeomorphism transforming one dynamical system into another. This justifies the use of mathematical methods to derive reduced phenomenological models that account for these crucial dynamical features in a more analytically and computationally tractable manner (Figure 3**A**). An example is the Wong-Wang model [90], which is derived under the adiabatic approximation [45] from the Brunel-Wang model [119] for a cortical network dominated by recurrent inhibition. Another approach to developing phenomenological models involves formulating hypotheses about the dynamical mechanisms–rather than the biophysical ones–that underlie specific features observed in the data [120]. This approach has proven successful, for example, in the modeling of epileptic seizures [97] or interictal spikes [121]. Given that the Virtual Epileptic Patient (VEP; [5, 8, 13]) is currently the most advanced application of the VBTs, we will devote the next section to an overview of NMMs for seizures, with a particular focus on the model of choice for the VEP, the Epileptor [97].

#### 3.1.3 Epileptor

The dimensionality reduction in the mesoscopic description is particularly justified in models of epileptic seizures, which are events characterized by a high degree of neuronal coordination. Among the models for seizures [87, 120, 122], those used for the region dynamics within large-scale personalized simulations mainly focus on the onset of seizures and on their propagation throughout the brain due to coupling among regions [121, 123–128]. The key features of most of these models are: the presence of two attractors–one for the interictal and the other the ictal state; a mechanism for transitions between these attractors; an excitability parameter characterizing how prone the system is to undergo a transitions; and a specific pair of bifurcations that destabilize the interictal/ictal states, influencing the overall dynamics of the model.

In more details, a fixed-point is used to represent the interictal (healthy), regime while a limit cycle is used for the ictal regime; in some cases, this is further reduced to another fixed point [127, 129]. Common dynamical mechanisms for seizure onset include [120, 130–132]: input- or noise-induced transitions among coexisting attractors (i.e., in the presence of bistability); and bifurcations that lead to the destabilization of the interictal state, when the bifurcation parameter is allowed to change due to noise or slow dynamics. Bistability is not required for bifurcations to occur, however, its presence can be exploited to produce hysteresis-loop bursting [133]. This provides a mechanism to autonomously produce seizures that does not rely on noise, thanks to feedback between fast and slow variables. This concept is central to the Epileptor model [97]. Another key characteristic of seizure models is the presence of a single parameter, generically referred to as excitability, that can be used to differentiate healthy and unhealthy brain regions and serves as the target for personalization at the NMM level, hence as a generative parameter in the inference process. The model can be further extended by including two neuronal subpopulations of epileptogenic and nonepileptogenic type, to make it capable of producing physiological oscillations in addition to the epileptiform activity [121]. Finally, different bifurcations leading to the destabilization of the interictal/ictal state exhibit specific signatures in the time series, for instance, the presence/absence of a baseline shift, or characteristic trends in the amplitude and frequency behaviors. These can significantly affect the dynamical properties of the model, including synchronizability or responsiveness to stimulation [97, 133, 134].

Despite the multi-factorial causes of epilepsy and the heterogeneity of its clinical manifestation, electrographic seizures display remarkably stereotyped features, consistent across species and even in small neuronal preparations [97]. Jirsa and colleagues studied these features, identified the signatures of specific bifurcations at seizure onset and offset, and proposed a taxonomy of seizures based on such bifurcations pairs [97, 110, 133]. They phenomenologically modeled the most common class in the Epileptor, incorporating systems that operate on three timescales: a fast subsystem, which displays the healthy fixed point and the limit cycle for fast ictal oscillations; a slow variable responsible for the transitions across attractors in the fast subsystem; and an intermediate subsystem, encoding both preictal spikes and spike-and-wave complexes. While the Epileptor is typically used in the slow variable regime, its parameters can be adjusted to accommodate both bistability and monostability [135], as well as transitions induced by input, noise, or slow variable–including mixed scenarios for seizure onset/offset [97, 136]– without altering the excitability parameter. Interestingly, the bifurcation diagram of the Epileptor shares core features with several biophysical models developed for epilepsy [88, 137, 138] (Figure 3**B**). Beyond modeling seizure onset and offset mechanisms, the Epileptor can also replicate phenomena such as status epilepticus and depolarization block [139].

However, patients also experience seizures that align with other classes of the taxonomy [97,134]. Some of these can be be accounted for by adjusting the parameters of the Epileptor [136], while these and others can be more generically modeled using unfolding theory [110, 140, 141]. This approach yields a simplified model with two fast variables and a slow one:

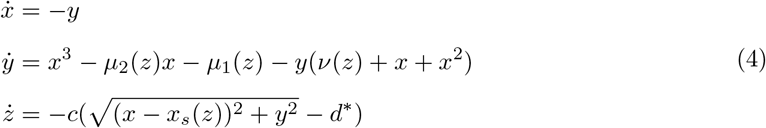

with *c* ≪ 1. The slow variable *z* pushes the fast subsystem (*x, y*) along a trajectory (*µ*_1_(*z*)), *µ*_2_(*z*), ν(*z*)) in parameter space, potentially crossing different bifurcations sequences, and generating seizures corresponding to specific classes (Figure 4, bottom). We refer to this model as generalized Epileptor.

#### 3.1.4 Data-driven models

Given the complexity of biological neural systems, designing a low-dimensional model aiming to capture the important features of neural dynamics is a difficult task, whether following more phenomenological or biophysical approach. Such task necessarily involves introducing some strong assumptions and approximations, whose applicability can be questioned. Modern machine learning, however, opens an alternative way; one that relies on experimental data and training techniques to learn the underlying dynamics. Such data-driven approaches can be understood as an extreme variant of phenomenological models, where the phenomena to be modeled are extracted from the experimental data automatically by appropriately designed training algorithms.

Data-driven models can be also used for model order reduction [142]. With this approach, a low-dimensional model is trained on data generated by a high-dimensional, computationally costly model– such as a model of large biologically realistic neuronal networks with single neuron resolution. The successfully trained low-dimensional model then captures the phase flows in state space generated by the high-dimensional model and thus serves as a first approximation of the latter’s emergent dynamics.

Range of methods for extracting the dynamics from data was proposed and used. Approaches based on linear state space models are theoretically well understood and can be efficiently trained [143], but are inherently limited in the nature of the dynamics they can represent. Even so, they provide a good baseline to evaluate more complex models of brain dynamics [144]. Recurrent neural networks (RNNs) are much more flexible, and are widely used for dynamical system reconstruction in neuroscience [145]. Their main building block is a state vector, supplemented by an update mechanism usually in form of some combination of matrix multiplications and element-wise nonlinearities. Popular variants include architectures designed for better performance on problems with long term dependencies such as Long Short-Term Memory (LSTM; [146]) or Gated Recurrent Unit (GRU) networks [147].

An approach related to RNNs is Neural Ordinary Differential Equations (NODEs; [148]), where the gradients of the loss function are calculated via adjoint method in continuous time, avoiding the use of memory-intensive backpropagation, and allowing the use of solvers with adaptive time steps. The disadvantages of RNNs include potentially difficult interpretation, since the trained system is not in the form of simple equations easily understood by humans. Better interpretability is the goal of the SINDy method [149], which approximates the system by a linear combination of pre-specified basis functions such as polynomials, and optimizes for sparsity in this representation. However, this approach faces challenges such as noise sensitivity, reliance on an appropriate function library, and balancing model sparsity with accuracy. Additionally, it struggles with highly nonlinear systems and encounters scalability issues in high-dimensional settings due to computational costs. Finally, sometimes a specific dynamical nature of a problem can be exploited to build a problem-specific form of a data-driven model, such as the network model of spreading epileptic seizures [150].

In the context of network-based brain modeling, the low-dimensional neural mass models can be trained either in isolation or embedded in the network. The former can be justified if the experimental data can be obtained from isolated tissue, or when trained on simulated data from larger model where the interregional coupling is either known or absent [151]. The trained neural mass models are then coupled together to form the whole-brain model using the methodology outlined in the next sections. In the latter case, the low-dimensional models are first coupled with the known structural network, and then their dynamics is trained jointly [152].

### 3.2 Connectome-based brain networks

Connectome-based models are a well-established approach for investigating functional neuroimaging modalities such as fMRI, MEG, and EEG [3, 49, 60, 101]. This network-based modeling incorporate individual structural brain imaging data [153], typically diffusion-weighted imaging data, to estimate the edge weights and lengths [154]. The structural connectivity imposes a constraint on the brain dynamics, allowing for the personalized simulation of the brain’s (ab)normal activities and their associated imaging recordings, potentially informing targeted interventions [5, 7, 13]. In the following subsections, we describe this approach in both low resolution (neural mass modeling) and high resolution (neural field approach).

#### 3.2.1 Networks with neural nodes at low resolution

At the coarser level of brain regions, the brain network dynamics are described as

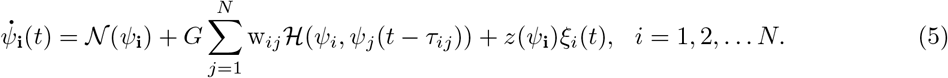

The evolution of local neural activity *ψ*_*i*_(*t*) in region *i* when uncoupled is driven by the inherent neuronal activity represented through the nonlinear function of the neuronal mass model 𝒩 (*ψ*_**i**_). Additionally, when constructing the brain network model, the connectome-based architecture dictates the strength and delay of the interactions between brain areas through the coupling function *ℋ* (*ψ*_*i*_, *ψ*_*j*_(*t* − τ_*ij*_)). Here, the axonal transmission introduces delays in the neuronal activity of distant regions, *τ*_*i,j*_, and the strength of this coupling is determined by the connectome weights w_*ij*_. These are typically estimated by tractography based on diffusion weighted MRI [155]. Finally, there is a noise term in the dynamics that can depend on neuronal activity through the function *z*, which multiplies a Gaussian with independent time series characterized by ⟨*ξ*_*i*_(*t*)⟩ = 0 and ⟨*ξ*_*i*_(*t*)*ξ*_*j*_(*t*^*′*^)⟩ = 2*Dδ*(*t* − *t*^*′*^)*δ*_*i,j*_.

Note that, in general, the working point of the system is determined by the interplay between the global coupling *G*, local (bifurcation) parameters, and the noise strength *D*. Time delays and weights constitute the space-time structure of connectivity, which is crucial in shaping its macroscopic activity [49,66,101,156], including the phase relationships between brain regions [157]. Their impact can be unified in the so-called normalization of the connectome [67] that allows graph-theoretical metrics to unveil structural affinities for spectrally-dependent activation patterns in the brain.

#### 3.2.2 Networks with neural fields at high resolution

To achieve a more spatially resolved activity of the brain, the driving Equation 5 requires to be modified to also include the interactions from the nearby tissue:

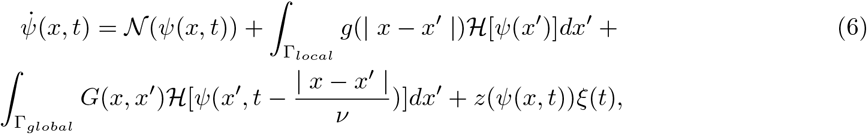

where, the neural activity *ψ*(*x, t*) at position *x* and time *t* is now also influenced by the short-range intracortical synaptic connections, which are instantaneous (i.e., *c* → ∞ in Equation 1). This assumes that neural populations across the cortical surface are connected to their neighbouring populations in a distance-dependent manner, as described by neural field theory [7, 28, 158]. To account for this in the model, the spatial resolution is increased. While in the brain network representation Equation 5, atlas-derived brain regions on the cortex could be collapsed to a single point (or node) in the network, the model now describes positions *x* as points on the unfolded cortical surface Γ_*local*_, as extracted from structural MRI. Typically local connection strength is assumed to decay with increasing distance and to be translationally invariant. Thus, a kernel is chosen and applied on the geodesic distances between points *x* and *x*^*′*^ on the cortex to obtain *g*(| *x* − *x*^*′*^ |). Common choices are Gaussian or exponential decay kernels, parameterized to limit the effective connection strengths to a few centimeters or millimeters.

In addition to the short-range, the long-range connectivity can now be represented as an integral over the spatial domain Γ_*global*_, extending across the whole brain. Assuming the heterogeneous two-point connection, 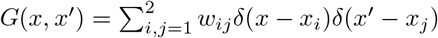, *i*≠ *j*, where *w*_*ij*_ ∈ ℝ represents the coupling strength through the heterogeneous connection [28, 158]. In the discretized version of the model, this increase in spatial resolution raises the number of positions *x*, from around 200 regions defined by the atlas to the number of vertices of the cortical surface mesh, which can commonly range up to 200 thousands. Thus, given a typical adult brain, a node of the connectome would approximate the neural activity of around 10 *cm*^2^ of the cortical surface, while with neural fields a single vertex represents around 1 *mm*^2^.

This increase in resolution results in a substantial increase in computational cost, potentially limiting the usability of the model, due to the formidable challenges it introduces for model inversion. Approximately 25000 iterations per second for the neural field simulation and 25 million iterations per second for the neural mass, representing a 3 orders of magnitude difference, were achieved on a state-of-the-art desktop GPU, the RTX 4090. Nevertheless, the pseudo-spectral approach [159] and reparameterizing the parameter space into spherical harmonic mode coefficients [160] can be employed to reduce the computational burden and facilitate the inference process. With this increase in resolution, certain empirically observed phenomena like cortical traveling waves [161] can be described. High spatial resolution is also essential for accurately modeling electric fields generated by neural sources or brain stimulation electrodes [5, 162].

### 3.3 Mapping neural sources to sensors

The neural activity *ψ*(*x*_*i*_, *t*), as simulated in Equation 5 and Equation 6, is not directly observable in real world experiments. Therefore a projection needs to be established to transform the simulated neural activity into empirically measurable quantities, such as BOLD fMRI, EEG, MEG or iEEG. For fMRI, the BOLD signal is a measure of deoxyhemoglobin in the tissue. Due to its paramagnetic properties, deoxyhemoglobin generates magnetic field gradients around blood vessels, leading to a reduction in the MR signal. The change in deoxyhemoglobin concentration is linked to neural activity through neurovascular coupling [163]. Multiple cellular and molecular mechanisms have been identfied which cause neural activity to induce vasodilation [164], including the release of neurotransmitters, ions, and metabolic byproducts that act on blood vessels to increase blood flow.

The typical BOLD signal due to a simple brief stimulus is described by the hemodynamic response function, which starts from baseline and typically peaks around 5 *sec* after stimulus begin, then decreases below baseline into an undershoot around 10 *sec* and, eventually returns to baseline after 20-30 *sec* [165]. The balloon model was introduced to explain the complex nonlinear response in the BOLD signal due to neural activity. When neural activity increases, blood flow rises to supply more oxygen to the active region. This causes the blood vessels to expand, much like a balloon, which in turn increases blood volume and alters the concentration of deoxyhemoglobin. The BOLD signal itself is a nonlinear response, depending on a complex interaction between blood flow, blood volume, and the amount of oxygen consumed [166]. The balloon model is given by a set of differential equations and was later extended to include a term for neural activity [165]. These equation can be used to transform simulated neural activity *ψ*(*x*_*i*_, *t*) into simulated BOLD fMRI. While the earlier balloon model assumed vasodilation of the venous system, more recent evidence suggests that dilation mainly occurs in the arterioles, leading to new hemodynamics models [167]. Similarly, improvements in the temporal and spatial resolution of fMRI imaging technology will require adaptations to hemodynamic models [168]. Furthermore, empirical evidence suggests that the hemodynamic response varies across brain regions and individuals, which may impact the the prediction of functional connectivity [169, 170]. Additionally, neurovascular coupling is often impaired in neurodegenerative disease, which may contribute to observed functional connectivity [171]. Similarly, the hemodynamic response changes with healthy aging [172]. Most studies using brain network modeling to predict functional connectivity so far have used a single balloon model across all brain regions, subjects, and conditions. This is probably a short coming of current studies. Future studies should investigate varying hemodynamic models to potentially enhance the accuracy of functional connectivity predictions.

For EEG, MEG and SEEG, a projection can be calculated by solving the electromagnetic forward problem. The forward model combines location and orientation of electromagnetic sources and sensors with a volume conductor model. The main source for the measurable electromagnetic field is the synchronous synaptic activity of groups of parallel-aligned cortical pyramidal neurons [173]. These synaptic transmembrane ion currents are approximated by a current dipole, positioned on and oriented orthogonally to the cortical surface, which can be extracted from a subjects structural MRI. The volume conductor is a three-dimensional model that describes the conductive properties of the extracellular tissue, including compartments such as the brain, cerebrospinal fluid, skull and skin. Next, the sensors location are aligned with the volume conductor and source model. Finally, the electric potential and magnetic induction at the sensors is calculated by solving Maxwell’s equation in its quasitatic approximation. In its simplest form, the volume conductor can be assumed to represent an unbounded and homogeneous medium, for which analytic solutions to Maxwell’s equation exist [174]. This assumption may be reasonable for intracortical sensors like SEEG, which are from any tissue boundaries and thus boundary effects may be neglected. For EEG and MEG, however, changes of conductivity impact the measured signal. In the simplest case, one- or three-layered spherical models, fitted to the head shape and sensors, have been applied. More advanced numerical approaches, such as boundary element methods (BEM) and finite element methods (FEM), use subject-specific geometries extracted from structural MRI for more accurate modeling [175, 176]. The BEM often considers three boundary surfaces: scalp-air, skull-scalp, and skull-brain. Conductivity within each compartmen is assumed to be homogenous and isotropic. In FEMs, the whole head volume is tesselated into tetrahedra, and a unique conductivity tensor can be specified for each tetrahedra separately. This allows to account for anistropic conductances in skull and white matter [177]. Further improvements can be achieved through biophysically realistic forward modeling, allowing for multiple dipoles, especially for sensors close to the neural sources, i.e., ECoG [178]. There are multiple openly available software suits which can be used to compute the forward model and to perform general analysis of empirical data, such as MNE [179], Brainstorm [180], FieldTrip [181], OpenMEEG [182].

## 4 Operating the Virtual Brain Twin: Technologies

In the following sections, we will detail the steps and technologies involved in operating the VBTs, starting with the preprocessing of structural data, construction of brain atlases, and coregistration to build individualized brain network models. This will be followed by an explanation of the simulation process and the inference techniques used to confront the VBTs with functional data.

### 4.1 Preprocessing, atlases and coregistration

Reconstructing virtual brains from neuroimaging data entails processing of multiple imaging modalities, including structural MRI (T1w and T2w images), diffusion weighted MRI, functional MRI, potentially CT scans, PET, and EEG/MEG/SEEG and additional modalities as applicable. The aim of the processing is to obtain the structural scaffold of a subject’s virtual brain, and processed functional data as a target for inference, against which dynamics of the model can be compared. The scientific field of image pre-processing is extensive, and we here point towards reviews from this field to give a broad overview. Preprocessing of diffusion weighted images involves a multitude of steps for denoising and artefact correction [183], followed by streamline tractography to estimate the white matter connections [155]. Subsequently streamlines are grouped according to some brain atlas [184] to obtain the structural connectome [185]. Furthermore, depending on the use case, post-SEEG implantation CT [186] or EEG-electrode MRI images [187] are aligned with the different imaging modalities to allow for an accurate electromagnetic forward model. Similarly, functional data like fMRI [188, 189] and EEG/MEG [190] are artefact corrected and aligned with the structural data. Often the voxel wise fMRI time series are averaged per brain region to compute the functional connectivity. For EEG/MEG, this can also be achieved following source estimation through an inverse model [191].

No gold standard pipeline has been agreed on within the neuroimaging community so far, and there is a large variety of possible steps to be included or not, which can affect the results, and consequently the input to virtual brain modeling. This variability was recently demonstrated in a meta-analysis, where 70 international independent teams analysed the same fMRI datasets with the pipeline of their choice. Interestingly, no two pipelines were identical, leading to sizeable variation in the statistical results obtained [192]. Similar studies have been conducted for tractography in diffusion weighted imaging datasets [193, 194], and another study on EEG data processing with the same intention is currently underway [195]. This highlights the fact that for well-grounded virtual brain modeling and inference, robust and well-tested image data preprocessing needs to be applied. To streamline the data preprocessing, different labs have packaged pipelines which take raw imaging data as input and give processed data as output. Each pipeline is differently adjustable and applicable to different quality of the input data. To name only a few examples: MICA pipe [196], fmriprep [197], nipreps (https://www.nipreps.org/), XCP-D pipe [198], MRtrix3 connectome [199], HCP minimal preprocessing pipeline [200],and a pipeline adjusted for virtual brain modeling on UK biobank imaging data [201].

### 4.2 Simulation

The simulation of a VBT (e.g., given by Equation 5) is an initial value problem for the differential equations representing neural activity, along with any forward models required. In the most general case, the equations are partial, stochastic and time-delayed,

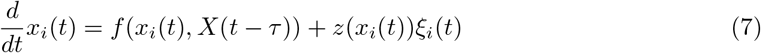

where *x*_*i*_ is the i’th state variable, *f* is the deterministic component of the model for the i’th variable, *X*(*t*−*τ*) are time delayed state variables, *z* is the diffusion component and *ξ*_*i*_ is a Brownian process. Note that the function *f*, which describes the dynamical properties of the system, encapsulates the local neural activity and the coupling function 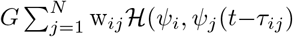 described in Equation 5. In practice, the partial derivatives are represented in a finite element fashion, thus not treated explicitly, additive diagonal Gaussian noise is used, time delays are fixed and follow a bounded unimodal distribution, and the equations are generally not stiff. While such equations can be solved with a variety of solvers, VBT solutions are usually obtained with an explicit stochastic Heun step, which is 2nd order in the deterministic component and 1st order in the stochastic component:

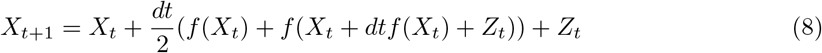

where 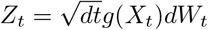 and *dW*_*t*_ ∼ *N*(0, 1).

In order to facilitate use by scientists, a Python package “The Virtual Brain (TVB)” has been written and maintained over the last decade, acting as a reference implementation [101, 202]. It is also available as open-source service on the cloud research platform EBRAINS (European Brain Research Infrastructures; ebrains.eu) [203]. The core library implements 29 different neural mass models, a handful of alternate numerical schemes, support for stimulation paradigms, along with several typical forward models for neuroimaging data, such as EEG, MEG, iEEG and fMRI. The fMRI solution is obtained either by solving the Balloon-Windkessel equations directly or via a linear approximation with a convolution. Some precalculation of so-called gain matrices for the EEG, MEG, iEEG forward models is provided, but we recommend using packages such as Brainstorm for this and importing the resulting gain matrix after for use in TVB.

The performance of Python, its NumPy array library and Numba JIT compiler in particular, enables pilot projects to design and test a specific model, but is insufficient for many “production” use cases, where the number of simulations to run for parameter sweep, simulation-based inference or Monte Carlo purposes is high. For these cases, tailored code is written either for CPU or GPU accelerators and supercomputing systems, usually involving adapting the layout of simulation work arrays for SIMD or SIMT parallelization, and high throughput schedulers such as Dask for distributing the work.

Several such implementations have been developed differing in the trade-offs between performance and productivity, as the translation of the mathematical formulation to the performance-oriented programming languages (such as C/C++) is time-consuming and error-prone process. The hand-written single-purpose implementations of a given simulation configuration such as [204] provide good performance, but are limited in extensibility. Two main approaches have been adopted to address this productivity bottleneck of the domain scientists. First, the structure of the reference simulator is replicated in the software components of the performance-oriented implementation to reduce the learning curve for the users adapting it to their needs, such as the *TVB C++* [205] backend of TVB. Second, the simulation is defined in a high-level configuration format such as XML, and the performance-level code is then generated based on the user input, effectively decoupling the domain experts from the implementation-related aspects of the simulation. This approach, taken by the `RateML` [206] also simplifies targeting different computing devices and integration of performance optimizations relevant for specific use-cases, such as the high-resolution models [207].

### 4.3 Model-based Inference

Inference is the method through which conclusions, decisions, or predictions are derived from initial premises, evidence, or existing data [208]. Broadly speaking, in statistics, inference is the process of discovering general principles about a set based on limited samples drawn randomly [209]. In machine learning (ML), it usually refers to using a trained model to generate predictions on unseen data [210]. In some cases it refers to discovering a statistical generative model in order to generate new observations that resemble the original dataset (generative adversarial network, transformers). In the context of VBT, the inference process relies on connectome based network model, as the data generative process, to simulate brain activity. The aims of inference are now to discover the most likely causes (within the scope of the model) of some unseen data and to predict new outcomes under some intervention in the model. By further integrating subject specific multimodal structural and functional data, the VBT becomes unique and personalized parameters are estimated to maximize the model’s predictive power, potentially informing clinical decision-making at the individual level. The evaluation of causal hypothesis also becomes possible in-silico, in scenarios [211–213] where testing in vivo, such as electrode implantation or the removal of epileptogenic zones in epileptic patients, would be prohibitive [5, 7, 13].

When estimating the subject-specific parameters that best explain the data, the inference process recasts to model inversion. The set of parameters to be estimated often includes regional parameters, such as control (bifurcation) parameters, along with a global scaling factor that defines the role of the structural connectivity matrix in revealing network effects. To incorporate uncertainty in the estimation and integrate background knowledge, such as clinical information [214,215], we use Bayesian inference, in which, all model parameters are treated as random variables, and their values vary according to their underlying probability distributions [216]. Bayesian inference is a principled method that updates the probability of a hypothesis based on new evidence and prior knowledge. In other words, Bayes’ rule combines and actualizes the prior information (background knowledge before seeing data) with the likelihood function (data-generating process) to form the posterior distribution (see Figure 6**A**). The posterior provides all the necessary information for evaluating accuracy and measuring out-of-sample predictive performance [217, 218]. Mathematically speaking, Bayes rule is defined as 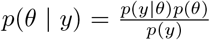 where, *p*(*θ* | *y*) is the posterior distribution of parameters given the observed data, *p*(*y* | *θ*) is the likelihood of data given parameter, and *p*(*θ*) is the prior distribution. The denominator *p*(*y*) = ∫ *p*(*y* | *θ*)*p*(*θ*) *dθ* is the marginal likelihood or evidence, which normalizes the posterior distribution. Methods differ in providing either samples from the posterior, or a tractable function that approximates the posterior function. Dynamic Causal Modeling (DCM; [219–221]) is a well-established approach in neuroscience that relies on approximation techniques, such as the variational free energy under the Laplace approximation (i.e., fixed-form Gaussian approximation to the posterior density), and it has been most widely applied to simple, linearly stable NMMs [222, 223]. VBTs typically use either adaptive Markov Chain Monte Carlo sampling, or training deep neural networks to learn distributions (without the need to specify a variational family or rely on factorization, Gaussianity or conjugacy assumption), while also incorporating nonlinearity [224–226].

**Figure 6:**
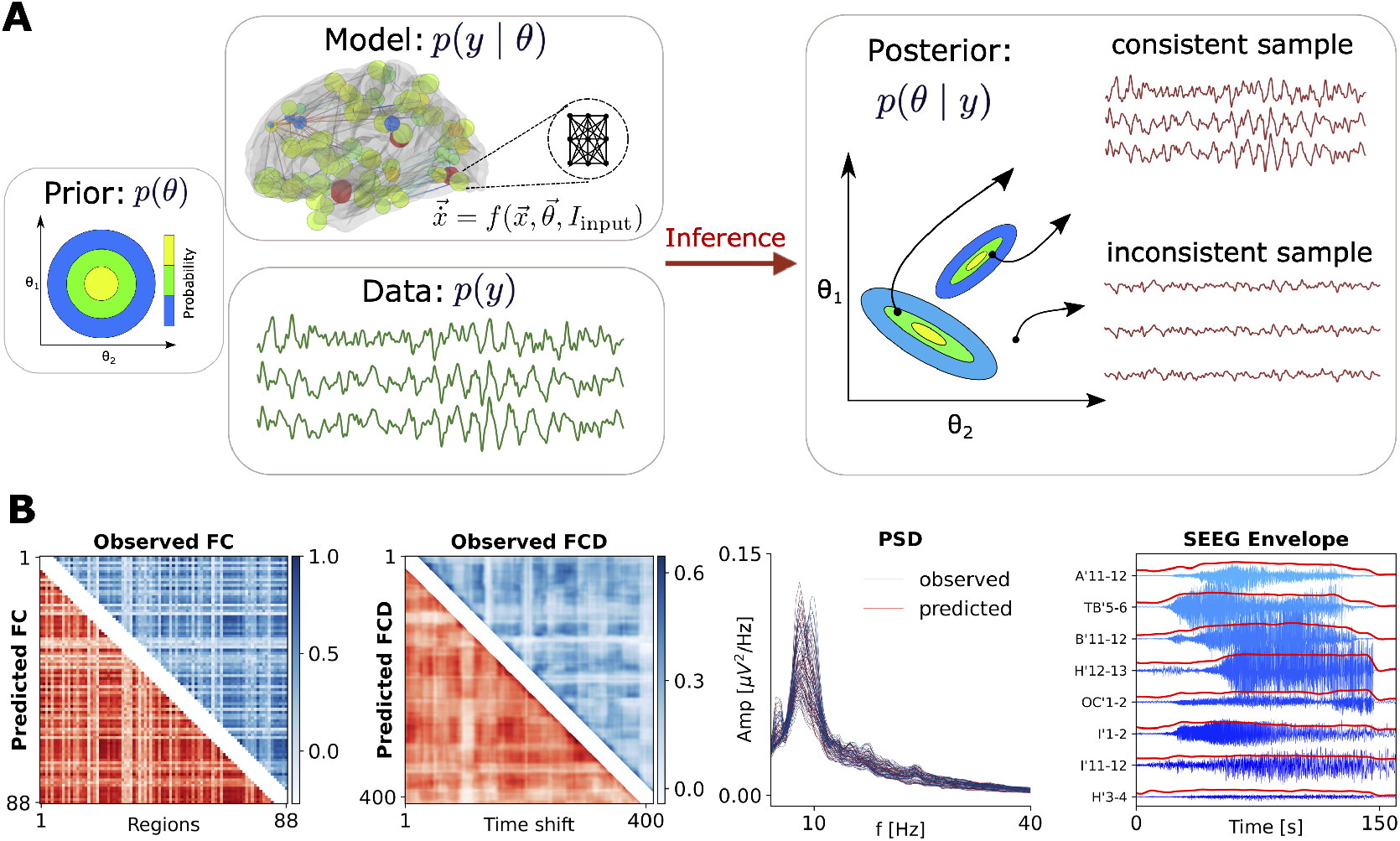
Bayesian inference in VBTs. (**A**) Based on Bayes’ theorem, background knowledge about control parameters in a VBT (expressed as a prior distribution, here in 2D), is combined with information from observed data (in the form of a likelihood function) to determine the posterior distribution. MCMC algorithms directly sample from the posterior distribution, while ML algorithms transforms a simple distribution (prior) into any complex target (posterior). Both methods inform us about degeneracy by providing all possible solutions in parameter space that best explain the data. (**B**) Examples of the observed and predicted data features in VBTs, such as functional connectivity (FC), functional connectivity dynamics (FCD), power spectral density (PSD), and SEEG envelope.

#### Markov Chain Monte Carlo

Markov Chain Monte Carlo (MCMC; [227]) sampling is a family of computational algorithms considered as the gold standard for unbiased sampling from the posterior distribution of model parameters. These algorithms evaluate consistency with empirical data through random simulations while preserving the correlation structure between parameters. Unlike point estimates such as maximum likelihood [44] or maximum a posteriori [228], Bayesian inference using MCMC provides the full posterior densities, allowing for a richer and more informative view of the parameter space, enhancing the quality of causal inference and decision-making. Adaptive and gradient-based MCMC sampling methods, such as the No-U-Turn Sampler [229], implemented in automatic probabilistic programming languages, like Stan [230], allow for accurate and efficient posterior estimation of parameters in VBTs. These techniques have been primarily opted for epilepsy as it involves complex dynamic brain activity patterns that require precise modeling [13,150,214,224,228,231]. The particular geometry of a realized likelihood function or posterior density function depends on the chosen parameterization of the model configuration space [232]. We introduced intricate reparameterization for these models to facilitate convergence and accurately capture and recognize their configurations [224, 231]. Accurate and efficient estimation of parameters in such models is crucial for understanding and predicting seizure events [160,233,234], which has driven the development of advanced MCMC methods in this field.

#### Simulation-based inference

Due to the high dimensionality of the parameter space and the data, as well as the nonlinear dynamics involved in neural mass models, MCMC methods often become computationally prohibitive. Of the alternatives, probabilistic ML algorithms, such as simulation-based inference (SBI; [235, 236]), can efficiently estimate the posterior distribution of parameters given only the low-dimensional data features (see Figure 6**B**). Tailored to Bayes’ rule, by using deep neural density estimators [237, 238] along with conducting random–but parallel simulations–, this class of probabilistic ML algorithms efficiently approximates the full posterior distribution of model parameters while reducing the computational challenges associated with calculating the likelihood function. Taking prior distribution 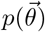 over the parameters 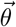, a limited number of *N* simulations are generated for training step 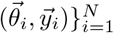, where 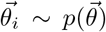 and 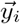 is the simulated data features given model parameters 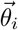 After training step, we are able to efficiently estimate the approximated posterior 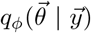 with learnable parameters *ϕ*, so that for the observed data 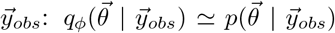 Importantly, SBI approach can be amortized at the subject-specific level, meaning that we do not need to retrain the model for new patient specific data [225, 226]. As a result, we can retrieve the posterior distribution of parameters in an order of seconds. This approach has demonstrated its capability in revealing pathological causes in brain disorders such as epilepsy [225], multiple sclerosis [16], Alzheimer’s disease [239], Parkinson’s disease [9], as well as in healthy aging [14] and focal interventions [240]. Note that identifying the set of low-dimensional data features that are informative of the control parameters for each case study is the main challenge in effectively applying SBI approach. Moreover, as an approximation technique, SBI may slightly overestimate the uncertainty or the relationships between parameters due to the limited number of simulations and the use of low-dimensional features. Nevertheless, this can be mitigated by using time-delay embedding techniques for data augmentation [151], or restructuring the model configuration space–such as through hierarchical reparameterization–to facilitate more efficient exploration of the posterior distribution [226].

## 5 Use cases of Virtual Brain Twin

In the following sections, we will demonstrate the use cases of the VBT, spanning the resting-state networks, healthy aging, multiple sclerosis, and epilepsy.

### 5.1 Resting state

Resting-state brain activity refers to the spontaneous neural processes that occur in the absence of specific tasks or external stimuli. Despite the absence of external demands, the brain’s activity remains highly structured both spatially and temporally (see Figure 7). This organization is marked by non-trivial correlations across distant brain regions, forming coherent patterns of activity known as functional connectivity (FC) [241, 242]. Over time, these patterns exhibit dynamic behavior, as strongly correlated functional communities are continuously formed and disrupted, giving rise to functional connectivity dynamics (FCD) [54, 116, 243, 244].

**Figure 7:**
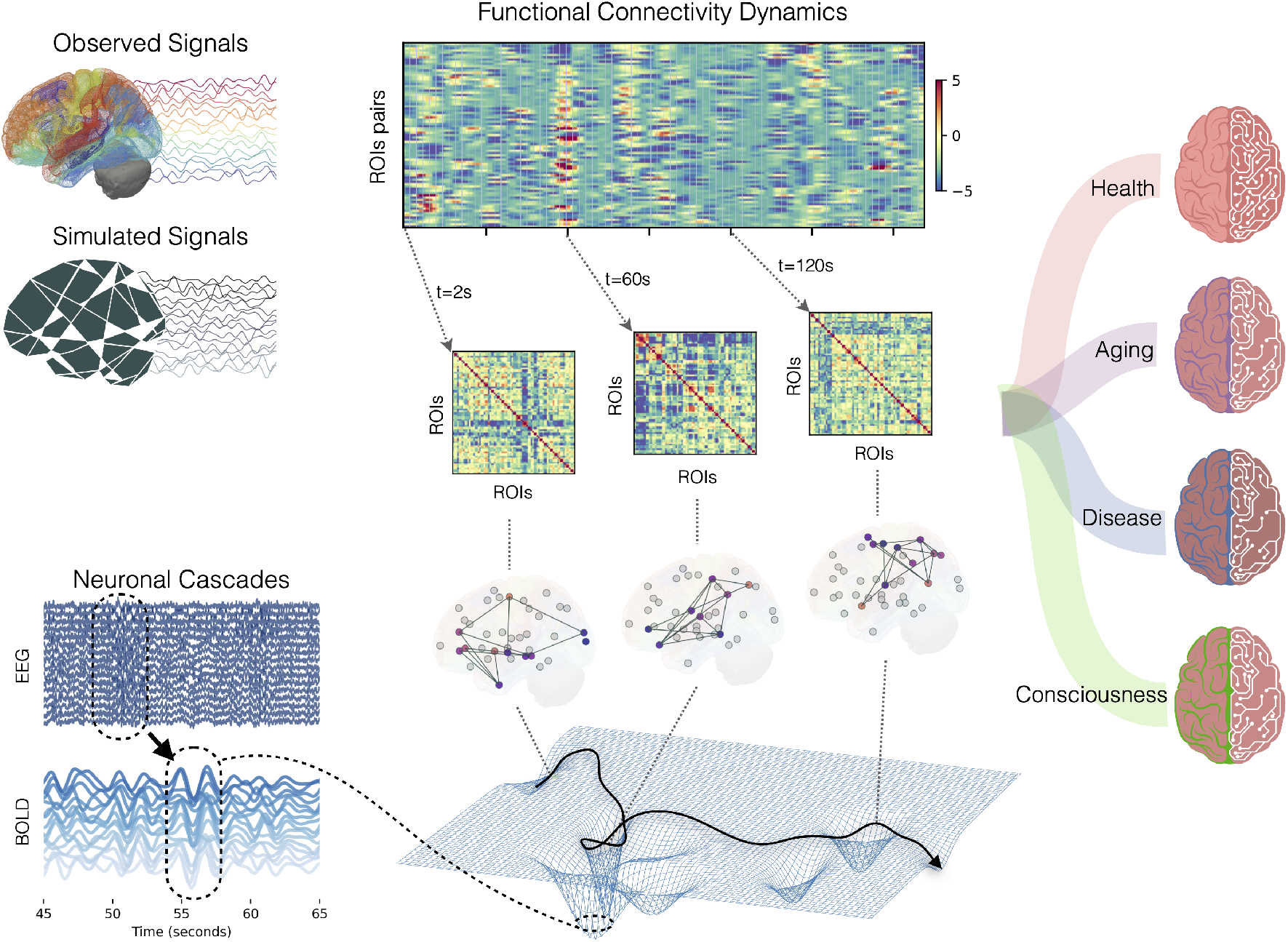
VBT of resting- and altered-states. Brain activity at rest is characterized by the dynamic formation and disruption of strongly correlated communities. Such characteristic dynamics are naturally simulated in VBT models which are built on empiric structural connectomes. One way to represent Functional Connectivity Dynamics (FCD) is via an edge-centric representation where, for each pair of regions of interest (ROIs), an edge co-activation time series is defined as the product between the standardized ROI-level signals. Averaging the co-activation signals (top raster plot) over time returns the Pearson’s correlation for each edge i.e., the static FC. Thus, at each time, the configuration of edge co-activations corresponds to an instantaneous realization of the FC (see example matrices). In time, the system explores various network configurations, which can be understood in dynamical system jargon as attractors on the Neural Manifold. In terms of neural signals, these correspond to high-amplitude fluctuations, or *neuronal cascades*, involving large-scale brain subnetworks, which generally underlie a slowdown of the system’s dynamics. FCD properties such as the variability of explored FC states (a.k.a., *fluidity*) or the richness of attractors dynamics (number, diversity, transition probabilities, etc.) are used as markers of health, disease, aging, and consciousness states.

Resting-state activity, often considered the brain’s default mode, serves as a baseline for cognitive processes [245]. Studying this baseline is clinically relevant, as alterations in resting-state FC have been linked to a range of neurological and psychiatric disorders, including Alzheimer’s disease [246, 247], schizophrenia [248], and depression [249]. Additionally, resting-state network dynamics provide biomarkers for aging [14, 250, 251] and various states of consciousness [252, 253] (see Figure 7).

While dynamic resting-state networks provide valuable insights into both healthy and pathological brain function, it is crucial to understand how deviations from healthy activity occur to gain mechanistic insights for diagnosis and prognosis in pathological conditions. Reorganization of whole-brain FC can be triggered by several factors, such as neuromodulatory changes [254], alterations in tissue excitability [255], and modifications in brain-gut communication [256], among others. However, without generative models of brain dynamics, discerning these mechanisms is challenging due to the brain’s emergent behavior, where studying individual parts in isolation cannot fully explain brain activity [257].

VBT offers a generative model that simulates brain dynamics resembling those measured empirically (Figure 7), enabling researchers to explore the underlying causes of FC reorganization. A key finding from these models is that a healthy brain operates near a critical-like regime [49, 116, 258]. Deviations from this regime are often observed during transitions in consciousness, aging, or disease states [259], with different parameter sets required to optimally capture such changes, ideally in a probabilistic manner (see subsection 4.3).

VBTs incorporate both the structural and dynamic aspects of brain function. Structural connectivity, which reflects the organization of white matter pathways, serves as the backbone for resting-state activity. Brain function simulated by VBTs relies on neural mass model equations (see subsection 3.1). Realistic resting-state BOLD fMRI activity can be modeled using meanfield equations (see subsection 3.1), with local parameters tuned close to instabilities, such as the Hopf model tuned near its bifurcation point [52]. However, while revealing the static dynamical core, this approach cannot identify its frequency-dependent aspects [67], which arise due to the interplay of the neuronal activity and the spatiotemporal profile of connectome’s weights and time-delays [49, 66]. A more comprehensive view emerges with the neural masses tuned to bistability and driven by noise to generate local fluctuations between low and high firing rates, resembling patterns observed during slow-wave sleep, anesthesia, and quiet wakefulness [117]. These models replicate the multistable attractor dynamics observed in real brain data [116]. The activation patterns produced by VBTs occupy specific structural modules and closely reflect the transient co-activation patterns identified in resting-state fMRI studies [117, 260, 261].

Although co-activation events are rare and deviate from baseline activity–raising concerns about their potential as artifacts rather than genuine neural phenomena–they remain significant contributors to FC. Brain models demonstrate that these events can naturally occur and should not be dismissed as artifacts. Further evidence from fMRI data across species supports the idea that transient co-activation patterns may serve as foundational components of FCD [262]. These transient functionally connected communities align with the brain settling into specific attractor states, while transitions between patterns indicate metastable dynamics [263–266]. According to synergetics principles [19, 20], when the brain enters an attractor state, a few macroscopic variables dominate, reducing the complexity of the system. This reduction allows brain activity to be modeled as a low-dimensional flow in state space [54, 267, 268]. In the resting state, structural connectivity introduces asymmetry between brain nodes, creating characteristic flow on the manifold, that governs the evolution of brain activity (see Figure 8). This manifold represents the space of possible functional configurations constrained by structural connectivity [47, 48, 269], offering insights into brain dynamics like traveling waves and cascade effects. Importantly, the manifold naturally constrains the inference process, enabling the sampling and estimation of a nuanced relationship between personalized parameters and large-scale brain dynamics, which is crucial for understanding clinical outcomes.

**Figure 8:**
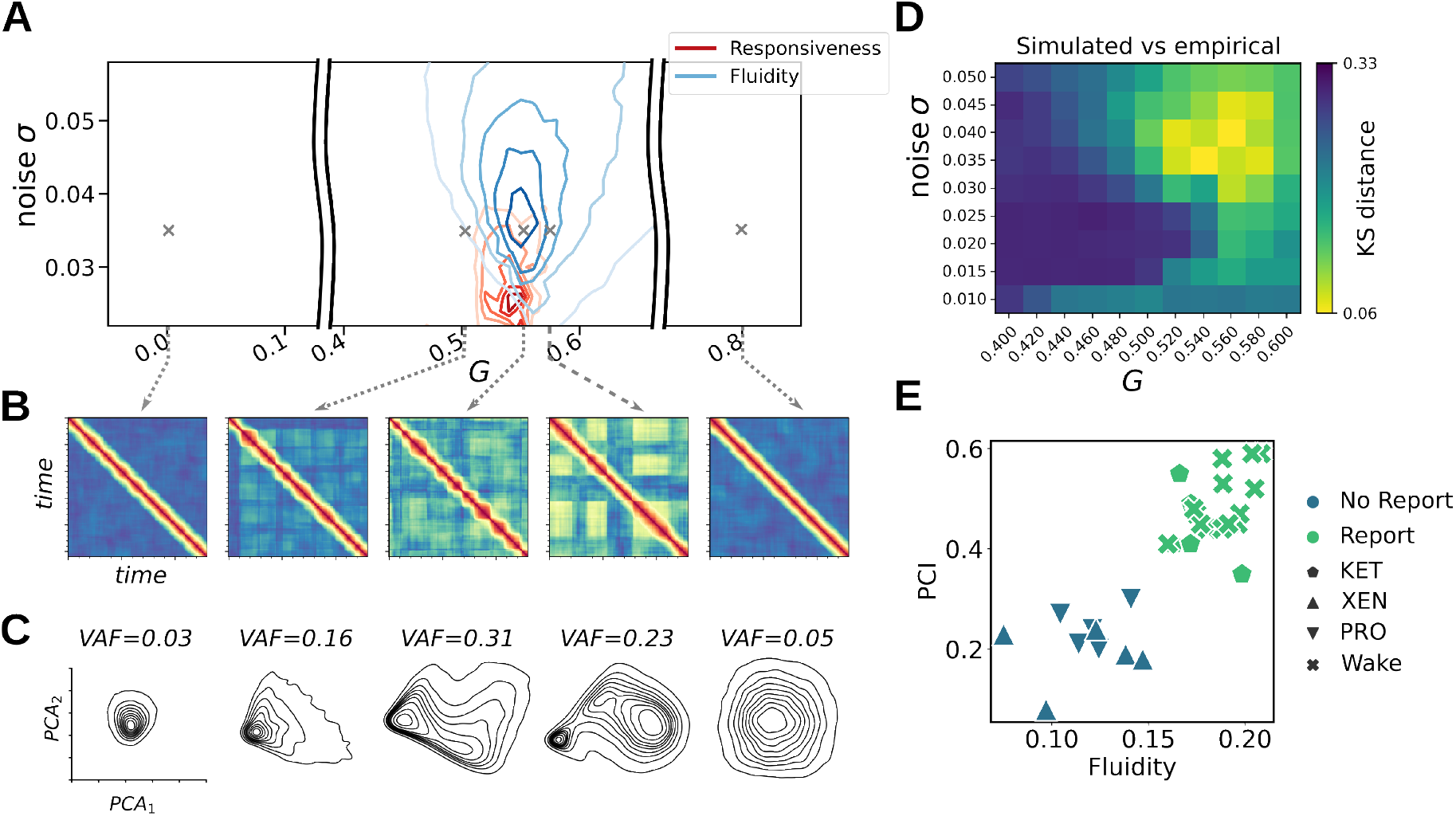
Structural Connectivity shapes the manifold of resting state activity. (**A**) The VBT exhibits the fluid dynamics in a narrow range of connectivity scaling *G*, where the local dynamics and the network influence are balanced and the symmetry across the nodes is broken (blue). The system in the working point also exhibits the highest responsiveness when the driving noise is decreased (red), suggesting the brain states with decreased responsiveness can be also tracked by fluidity of the spontaneous activity. (**B**) The fluid dynamics as reflected in the Functional Connectivity Dynamics (FCD) of the simulated BOLD signal. (**C**) The dimensionality of the dynamics decreases in the working point, here shown in the projection of the model state variable time series into the first two PCA components, and assessed by the Variance Accounted For (VAF) by these two components. (**D**) The simulated activity is most similar to the empirical data (Kolmogorov-Smirnov distance between the FCD matrices) in the working point. (**E**) The prediction from the VBT regarding responsiveness was validated in the empirical data in subjects under anesthesia (Propofol, Xenon, Ketamine). The fluidity of the spontaneous activity performed as well as the perturbation complexity index in separating the cases of conscious report. Figure was adapted from our previous studies [48, 271].

VBTs have revealed the role of the neuromodulatory system in shaping brain dynamics by regulating the attractor landscape [254]. The organization of resting-state activity also underpins the brain’s capacity to respond to external stimuli. Using VBT models, it has been shown that the complexity of stimulus-evoked responses, a reliable marker of consciousness [270], is tied to the same parametric regime in which spontaneous activity remains fluid i.e., flexibly switches between attractors [271] (see Figure 8). Fluidity measures applied to resting wakefulness versus anesthesia demonstrate a comparable ability to assess consciousness as traditional stimulation paradigms.

A striking example of the brain’s interconnectedness is the widespread effect of local manipulations, such as focal lesions or optogenetic neuronal silencing, on resting-state brain organization [272–275]. VBTs confirm these findings, linking the observed reconfiguration to either structural lesions [276] or changes in local neuronal excitability [277]. In sum, VBTs offer powerful tools for formulating causal hypotheses and interpreting experimental data. By providing mechanistic insights into brain dynamics, they facilitate a deeper understanding of both healthy and pathological states of brain function.

### 5.2 Healthy aging

Aging of the brain is well described, both structurally and functionally [278,279]. On the structural side, it leads to a substantial overall decrease in the number of streamlines [278], which mostly affects inter-hemispheric links that decrease with advancing age [14, 251]. On the functional side, applying graph theory metrics reveals age-related increases in between- and decreases in within-network resting-state FC [280]. However, FCD and higher order connectivities are more predictive of the brain age than the static FC [251], with age-related slowing down in FCD becoming evident when it is characterized as a random walk [250].

The observed patterns between structure and function have been statistically linked [281], including studies using TVB [282]. TVB was also used to demonstrate a link between the decrease in complexity of the brain’s function with aging and its structural changes, represented as long-range pruning [283]. Similarly, demyelination and white matter atrophy, especially the decline of the long interhemispheric tracts, have been linked to changes in FCD and metaconectivity at the individual level, with a key role of dynamical compensation through the increased global coupling [251]. However, the individual-level variability of structure and function links to cognitive decline, has only recently been demonstrated [14].

Lavanga and colleagues built a Virtual Aging Brain (VAB; [14]) framework to infer the causal link between structural connectivity (SC) and functional architecture, and how this impacts the consequent cognitive decline in aging of the 649 healthy subjects in the range of 55-85 years from the 1000 Brains cohort [284], as schematically shown in Figure 9**A**. Based on the analysis of the personalized data and earlier works [251], interhemispheric degradation of SC was identified as a primary aging mechanism, Figure 9**B**. Testing this in-silico, they reproduced the process of functional dedifferentiation during aging, Figure 9**C**. In particular, the study hypothesized that experimentally decreasing the interhemispheric SC (presented by the *α* mask parameter) would mimic a major mechanism of the structural aging process in the brain (i.e., “virtual aging”). This change would consequently be associated with reductions in the interhemispheric fluidity and the homotopic FC, as these two functional patterns demonstrate the highest sensitivity to aging. While setting up the working point of the virtual brains, the authors hypothesized that the brain attempts to maximize its fluidity (with FCD variance serving as a proxy) based on previous works that identified the FCD variance as an indicator of brain fluidity, which has been linked to cognitive flexibility [117, 250, 285]. Based on this criterion, the VAB framework presented a significantly higher coupling between SC and FCD, indicated by a higher global network parameter *G*, with increased age, but also with stronger SC deterioration (corrected for sex and education), as shown in Figure 9**C**. Hence, this demonstrates that the global modulation, which was hypothesized to be linked with neuromodulation–particularly the dopaminergic processes [286]–plays a compensatory role in preserving cognitive capabilities, and it increases with age and with SC deterioration.

**Figure 9:**
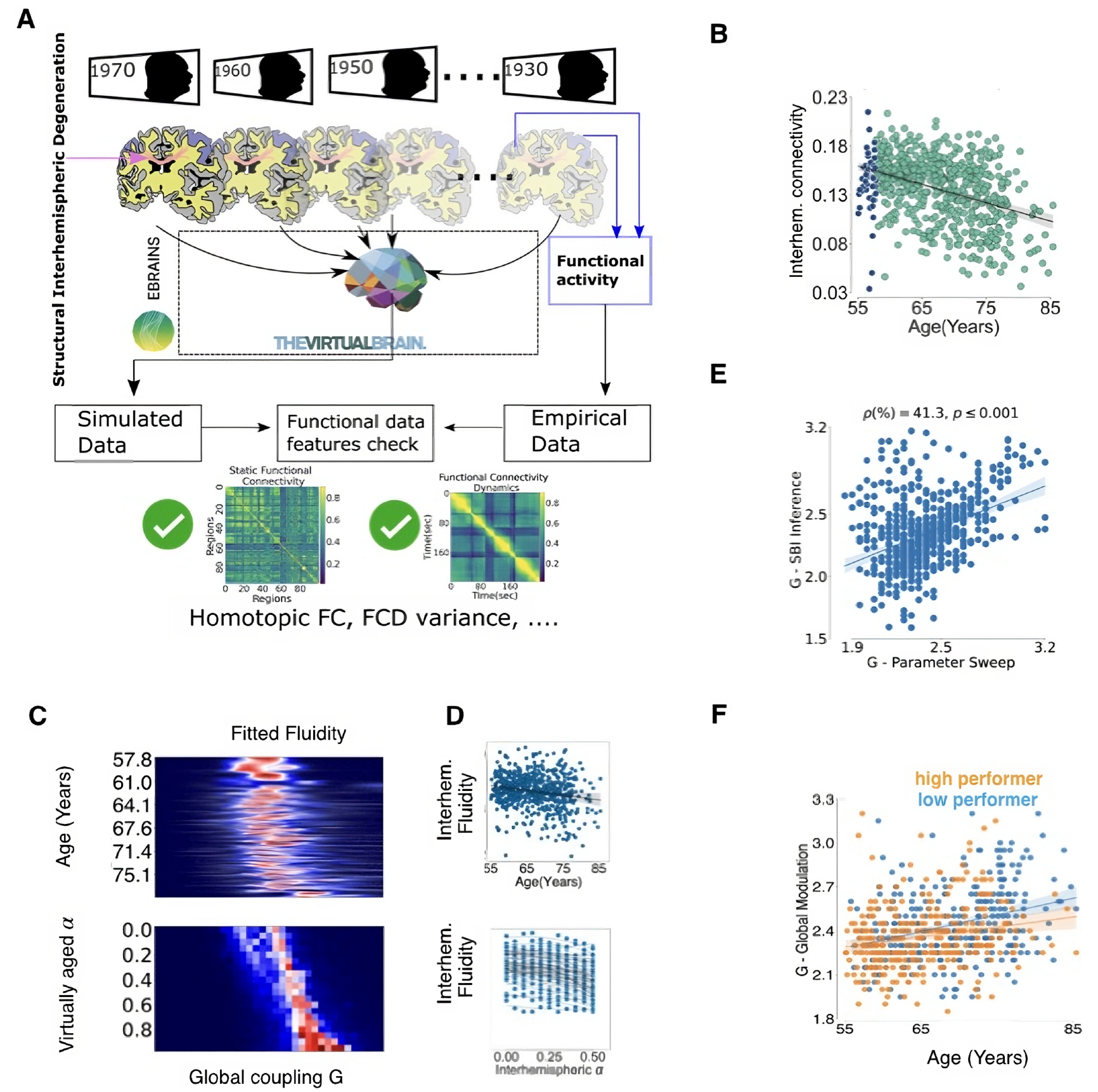
Virtual Aging Brain framework. (**A**) Virtual brains are constructed using TVB from a cohort of healthy subjects aged 55 to 85 years. The fMRI BOLD signals are simulated, and selected data features (homotopic functional connectivity and FCD variance) are compared with empirical data. (**B**) Interhemispheric SC decreases in the empirical data (*ρ* = − 0.453, *p <* 0.001. The 50 youngest subjects, represented in blue, are later virtually aged. (**C**) Heatmap of the simulated FCD variance and its trend with age (top) and virtually aged through degraded interhemispheric connectivity (bottom). (**D**) Interhemispheric FCD variance as a metric of fluidity for the virtual subjects (top) and the virtually aged subjects (bottom). (**E**) The correlation between *G* modulation index obtained with Bayesian SBI and the *G* obtained by parameter sweep. (**F**) The aging *G* trend split by concept shifting: low performers (blue) and high performers (orange) (Fisher’s-Z: *p* = 0.046, *ρ* corrected for sex and education). Figure was adapted from our previous study [14].

The Bayesian SBI [225, 226] also confirms the increase in SC neuromodulation with aging on an individual basis, as shown in Figure 9**D**, and retrieves global modulation with the same age related declines in FC and FCD features. This provides a causal validation of the VAB pipeline and suggests that higher model evidence corresponds to increased *G* values with age, indicating a shift in the brain’s optimal working point as it ages. Given that the age-related increase in SCFC coupling is characterized by functional differentiation [287], the slightly stronger age-related increase observed in *G* in low-performing individuals, Figure 9**E**, might additionally reflect an acceleration of these processes. In addition to being applied in a task-free paradigm, this is direct evidence of scaffolding and dedifferentiation in aging [287], leading to adverse effects of cognitive decline as demonstrated within a subject-specific causal inference framework in a large cohort.

### 5.3 Multiple sclerosis

Multiple sclerosis (MS) is a chronic, autoimmune, and degenerative disease of the central nervous system, affecting approximately 2.8 million people worldwide in 2020 [288]. The disease involves the immune system attacking the myelin sheath, which insulates nerve fibers (axons) and supports efficient electrical signal transmission, leading to a range of motor and cognitive symptoms [288]. VBTs can be particularly valuable for stratifying patients due to the heterogeneous nature of MS [16] and for predicting the effects of therapeutic changes, such as switching treatments. In the case of MS, the pathophysiological mechanism directly affects the large-scale connectivity, which can be promptly accommodated in models available in the VBT. Today, clinical monitoring relies heavily on structural lesions, that appear as a result of pathological activities. However, structural lesions correlate poorly with clinical impairment, i.e., the clinical-radiological paradox [289]. The expanding array of treatment options and the availability of extensive multimodal datasets make it necessary to monitor pathological activities before structural lesions appear.

Current predictive models for MS have focused on individual responses to disease-modifying therapies, often employing generalized linear models [290]. These studies aim to forecast clinical outcomes based on large, multidimensional datasets but do not provide a direct mechanistic explanation for developing patient-specific disabilities. Recently, a novel model has been proposed to infer conduction velocities from large-scale brain data, the so-called Virtual Multiple Sclerosis Patient (VMSP; [16]). This probabilistic approach is based on the notion that symptoms are linked to reduced conduction velocities, which are challenging to measure directly across the brain. Typically, structural lesions are assessed, via magnetic resonance, to determine damage accumulation and therapeutic responsiveness [291], but VBTs offer the potential to infer conduction velocities more directly.

Patients with MS exhibit larger functional delays throughout the brain compared to healthy individuals, particularly in regions with structural lesions [292]. As explained, changes in myelination can disrupt the timing of interactions between brain regions, leading to symptoms. In this context, the time delay represents the quantity to be inferred, as it is most directly related to the pathophysiological mechanisms. Recently, leveraging a multimodal dataset with MEG and tractography, it was shown that it is possible to infer the average conduction delays in individual patients starting from the spectral properties measured from source-reconstructed MEG data [16]. To this end, we constrain the Jirsa-Haken equation, given by Equation 1, to operate the neural masses locally in the oscillatory regime, which forces an emphasis on the full space-time structure of the connectome, notable the time delays via signal propagation. In such reduced form, this can be written using Stuart-Landau oscillators (known as the normal form of a Hopf bifurcation, see Table 1) coupled through to the empirical patient-specific connectomes as in:

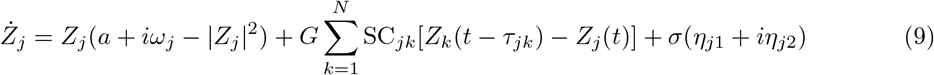

where *Z* is a complex variable, and Re[*Z*(*t*)] is the corresponding time series. Each region has a natural frequency of 40 Hz (*ω*_*j*_ = *ω*_0_ = 2*π* · 40 *rad/s*). Note that coupling between regions (with strength of *SC*, and scaled by *G*) accounts for finite conduction times, which are estimated by dividing the Euclidean distances between nodes by an average conduction velocity *τ*_*jk*_ = *d*_*jk*_*/*ν (see Equation 6, and Equation 5).

The low-dimensional representations of data were used to obtain the expected power spectra given specific delays (see Figure 10**A, B**), as they effectively capture the collective frequency emerging from synchronization mechanisms [16, 17]. For weak couplings and large delays, the network resists synchronization, and oscillators drift close to their natural frequencies (40 Hz), whereas stronger coupling is associated with the emergence of slower and lower amplitude oscillations [66, 293]. Assuming intermediate coupling (not sufficiently strong to stabilize full synchronization), summary statistics of PSD were used to train a class of deep neural density estimators within the framework of simulation-based inference (see subsection 4.3). Furthermore, the most likely individual average delays demonstrated to have predictive power over the clinical disability (Figure 10**C**). However, MS poses a very specific challenge, in that the damaged edges are specific to each patient. In other words, not all edges are damaged in every subject, and the damaged connections vary between patients. On the one hand, limiting oneself to average delays provokes a relevant loss of information about the patient-specific topography of the lesions. On the other hand, inferring the delays for thousands of edges at once becomes computationally prohibitive, due to the high dimensionality of the parameter space and the ubiquitous degeneracy, which leads to waste of computational resources. To overcome this issue, a novel approach was proposed, starting from a linear formulation of the relationship between the conduction delays and the amount of structural damage in each white matter tract [17]. Dependent upon a single control parameter, this function translates the intensity of empirical lesions into edge-specific conduction delays, which in turn lead to shifts in the power spectra (Figure 10**D**). The estimated parameters (obtained via a neural density estimator) were cross-sectionally predictive of individual clinical disability [16, 17]. This study represents an initial exploration showcasing the location-specific impact of myelin lesions on conduction delays, thereby advancing the customization of VBTs for individuals with MS.

**Figure 10:**
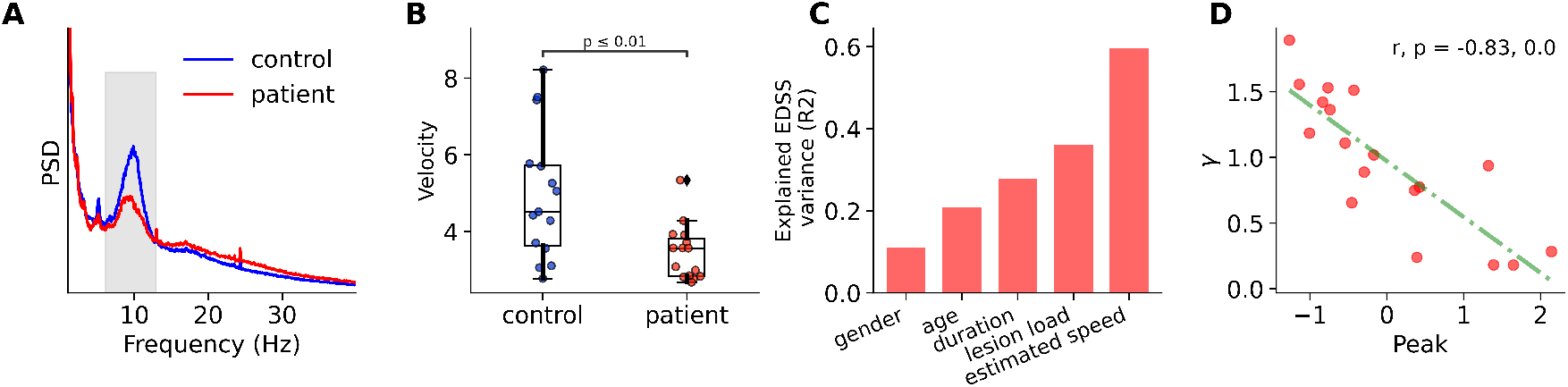
Virtual Multiple Sclerosis framework. (**A**) Median power spectral density (PSD) plotted for controls (blue) and MS patients (red). Shaded region represents the alpha band (8–13 Hz). (**B**) The inferred velocity parameter significantly decreases in the control group relative to the patient group. (**C**) Variance explained by the additive model including five variables (i.e., gender, age, disease duration, lesion load, and estimated speed). Adding the estimated conduction velocities significantly increased the predictive power. (**D**) Correlation between inferred parameter *γ* (translating lesion into edge-specific conduction delays) and the amplitude of alpha-peak in MEG recordings. Figure was generated using methods and data from our previous study [16, 17].

### 5.4 Epilepsy

Epilepsy affects around 50 million people worldwide and can cause to long-term disability. The burden of epilepsy arises from several factors, including a high prevalence of drug resistance (around 30% of cases) and comorbidities such as depression and anxiety [5]. Seizure generation and propagation involve large-scale neuronal networks, and the complex spatiotemporal dynamics in recorded data cannot be easily explained by the traditional model of a single seizure focus triggering activity that spreads to non-epileptogenic brain regions [127, 128].

The first practical use case of VBTs was developed for epilepsy, known as the Virtual Epileptic Patient (VEP; [8]). The initial concept for constructing personalized whole-brain network models in epilepsy was introduced previously [8, 224], where the key building blocks were detailed. This first application focused on estimating the epileptogenic zone networks (EZN). The VEP workflow for estimating EZNs was evaluated using data from 53 patients with drug-resistant focal epilepsy [13, 294]. VEP is currently being evaluated in an ongoing clinical trial (EPINOV NCT03643016) with an enrollment of 356 prospective patients. The VEP is now also being used to predict surgical outcomes through virtual surgeries [13] and to explore electrical stimulation techniques such as SEEG and temporal interference stimulation [295].

Here, we briefly describe the modules of the VEP workflow, which can be considered standard construction patterns for VBTs in both healthy and diseased brains. The VEP extracts individual brain anatomy from T1-weighted MRI (T1-MRI) and defines 162 brain regions according to the VEP atlas [296]. Tractography is performed on diffusion-weighted MRI (DW-MRI) data to estimate the length and density of white matter tracts (subsection 4.1). These tracts are then grouped according to the regions defined by the atlas, allowing for the derivation of a structural connectivity matrix that specifies the connection strength between brain regions. The post-SEEG implantation CT scan is used to determine the exact locations of the SEEG electrodes and to construct the source-to-sensor map using the gain matrix (subsection 3.3). The gain matrix maps the simulated neural source activity to the corresponding SEEG signals. Neural mass models at each source location are connected via the connectivity matrix, and neural source activity is then simulated (subsection 3.1). Finally, Bayesian inference methods are employed to estimate patient-specific parameters of the model based on features extracted from SEEG signals and priors derived from additional knowledge, such as SEEG data analysis or clinical hypotheses (subsection 4.3). The output is the suggested EZN, which can be represented in a distribution table and a heatmap overlaid on the T1-MRI data.

Figure 11 presents a patient example to illustrate how the VEP works. This 29-year-old right-handed female patient was initially diagnosed with left frontal epilepsy (Figure 11**A, B**). Adaptive optimization and MCMC sampling algorithms were used to estimate the key parameters of the VEP models based on the data features from SEEG seizure recordings. These key parameters, including the excitability of each region and the global connectivity coupling, determined the epileptogenicity values for each region he VEP identified the EZN based on the distribution of epileptogenicity values (Figure 11**C**). Based on the clinical hypothesis, the EZN was thought to involve the left insula gyri brevi and the left orbitofrontal cortex (Figure 11**D**, shown in yellow). The patient underwent resective surgery targeting the clinically defined EZN, which resulted in a reduction in seizure frequency but did not achieve seizure freedom. According to the VEP, the EZN included not only the left insula gyri brevi and the left orbitofrontal cortex but also the left inferior frontal sulcus (Figure 11**D**, shown in red). The left inferior frontal sulcus was an additional region identified by the VEP. Based on this finding, and after reviewing all related patient data, the patient underwent a second surgery. The patient has been seizure-free for over two years.

**Figure 11:**
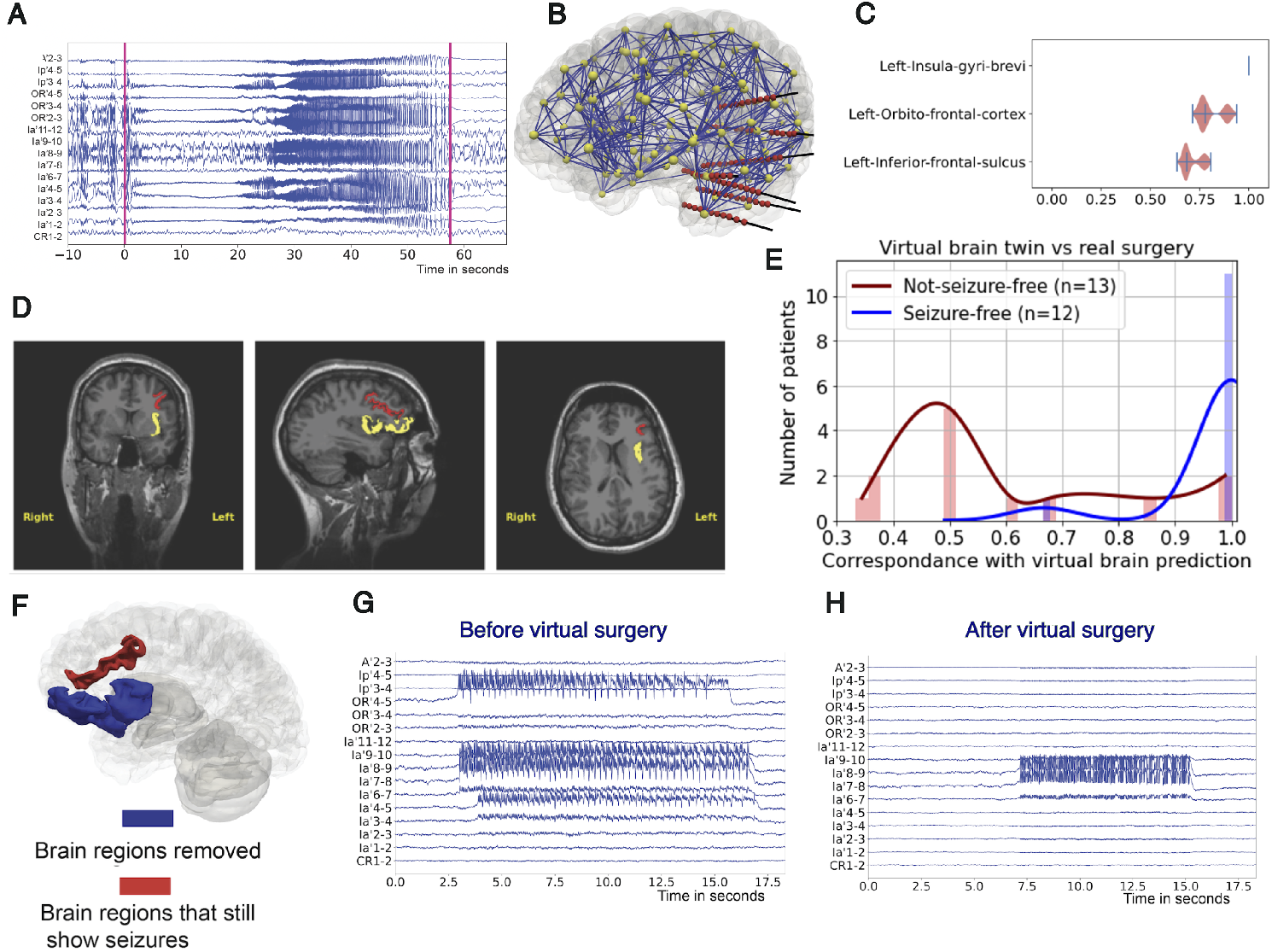
Virtual Epileptic Patient framework. (**A**) Selected SEEG recordings from a 29-year-old female patient. (**B**) Personalized whole-brain models: brain region nodes are shown in yellow, connectivity links in blue, and SEEG electrodes in red. (**C**) Posterior distribution of the EV (where higher values indicate a higher probability of seizure) for three selected regions obtained from the MCMC sampling pipeline. (**D**) Heatmap of regions identified exclusively by VEP (in red) and overlapping regions with the clinical hypothesis (in yellow), displayed on a preoperative T1-MRI. (**E**) Precision of VEP compared to postoperative MRI in 12 seizure-free patients and 13 non–seizure-free patients. (**F**) The left insula gyri brevi and left orbitofrontal cortex (in blue) were virtually resected, while the left inferior frontal sulcus (in red) continued to show seizure activity. (**G**) Selected simulated SEEG time series using the estimated EZN by VEP pipeline. (**H**) Selected simulated SEEG time series after virtual surgery. Figure was adapted from our previous study [13].

The VEP workflow was retrospectively evaluated using data from 53 patients with drug-resistant focal epilepsy [13]. VEPs accurately reproduced the clinically defined EZNs with a precision of 0.6, where the physical distance between epileptogenic regions identified by VEP and the clinically defined EZNs was small [13]. When comparing the VEP results with the resected brain regions, precision was used as a performance metric, highlighting the importance of minimizing false positives. In seizure-free patients, it is reasonable to assume that the EZN was completely removed, and thus any false-positive estimate is likely to be truly outside the EZN. Conversely, in non–seizure-free patients, a false-positive estimate is likely to correspond to non-resected epileptogenic regions, which may contribute to the persistence of seizures. The VEP demonstrated a very high precision (mean, 0.972) for seizure-free patients (Figure 11**E**). In contrast, the VEP results for the non–seizure-free group showed a significant decrease in precision (mean: 0.593) [5, 13], suggesting that there may be potential to better exploit the predictive power of the VEP. It should be noted that, in some patients, the very large extent of resection also contributed to the high precision value to some degree.

VBTs enable the simulation of brain activity data under various conditions, such as different stimulation protocols, seizure onset scenarios, or interictal states. As shown in Figure 12, once the VBT is constructed from the patient’s specific data, we can simulate brain activity and map it to SEEG signals, enabling a direct comparison with the patient’s empirical SEEG signals. The simulated signals from the VBT, generated with subject-specific control parameters, can provide a ground truth for detailed data analysis studies, such as evaluating algorithms and methods for estimating the EZNs [13] or testing EEG and MEG source localization algorithms [298, 299].

**Figure 12:**
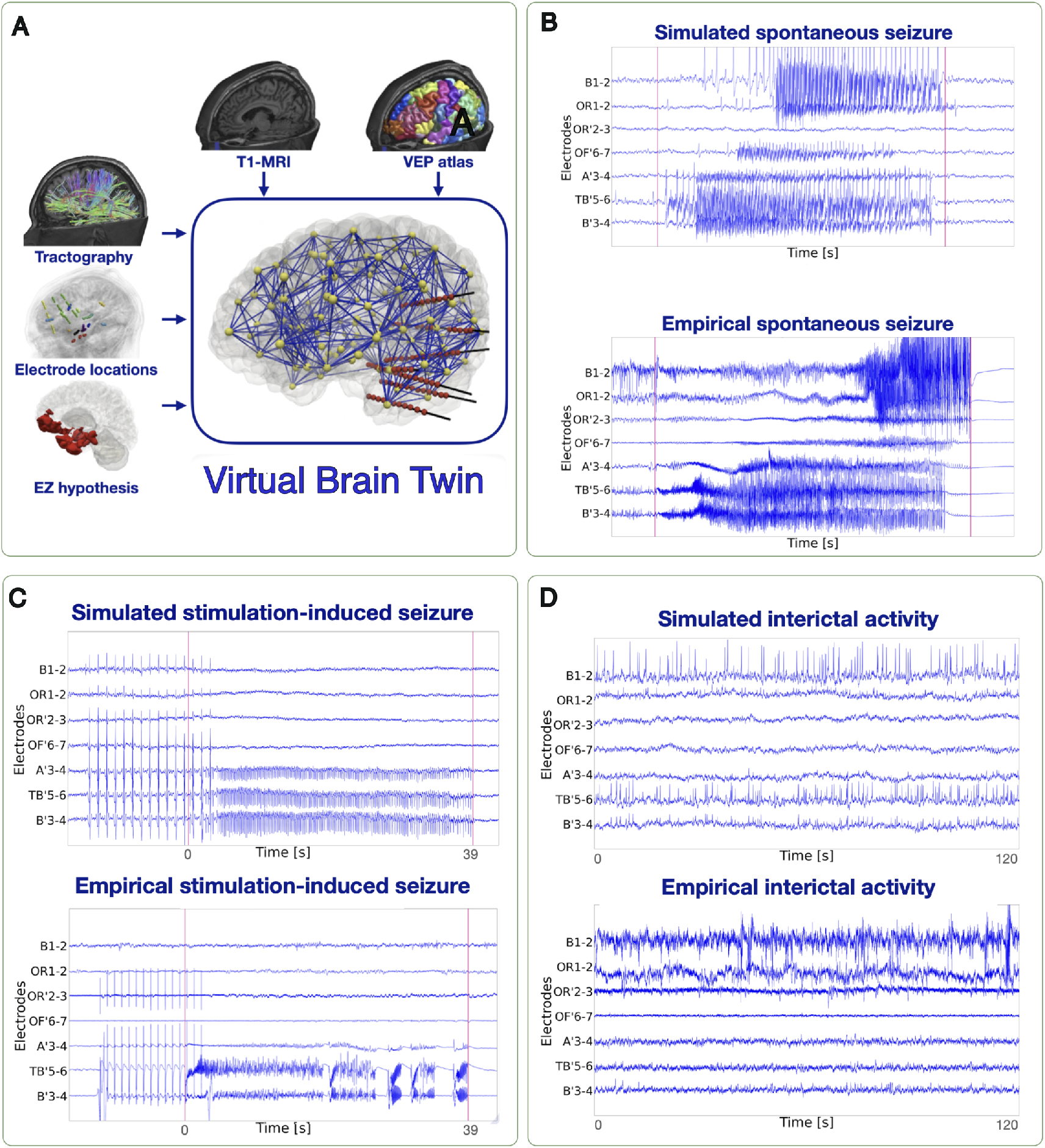
SEEG time series in the VBT of epilepsy. (**A**) The VBT was constructed using the patient’s specific data. For the same patient, the time series of simulated SEEG signals (top) and empirical SEEG signals (bottom) are shown for three different conditions: (**B**) Spontaneous seizures, (**C**) Stimulation-induced seizures, and (**D**) Interictal activity. Figure was adapted from [297].

Another practical use of VBTs is providing virtual surgery results for patients with drug-resistant epilepsy. Epilepsy surgery is the only curative option to stop seizures in individuals with drug-resistant focal epilepsy [5]. However, the failure of surgery can result from several factors inherent to epilepsy (e.g., involvement of eloquent cortices or extended networks), the surgical procedure (e.g., insufficient resection), or SEEG assessment (e.g., inadequate investigation, sampling limitations, or challenges in interpreting findings). A follow-up SEEG assessment after unsuccessful epilepsy surgery may reveal an additional epileptogenic area near the original zone or in distant regions. The VBT has the potential to improve postsurgical outcomes by enhancing the prediction of the extent of the epileptogenic zone, the behavior of non-operated regions, and the potential effectiveness of surgery [13, 300, 301]. For instance, VBT can provide information on brain regions not sampled by SEEG and be used for model testing under conditions different from those used in the model’s construction, such as simulating different clinical hypotheses for ictal and interictal states and testing surgical strategies (Figure 11**F-H**). These applications highlight a key advantage of VBTs over model-free approaches, which are limited to the regions explored by SEEG [5].

## 6 Outlook and discussion

Virtual Brain Twins (VBTs) offer a promising avenue for advancing our understanding and treatment of neurological and neurodegenerative disorders. By building brain network models that integrate individual structural and functional data, VBTs enable the (low- and high-resolution) simulations of brain dynamics, which, through Bayesian inference of personalized parameters and their associated uncertainty, provide insights into causal mechanisms. In addition to realistically simulating healthy resting state and normal aging, VBTs allow for in-silico testing of interventions when in-vivo experiments are prohibitive (e.g., in epilepsy), enabling more tailored and precise treatment strategies, thus extending their utility to other brain disorders.

Parkinson’s disease (PD) is the second most prevalent neurodegenerative disorder, characterized by motor symptoms such as tremors, rigidity, and bradykinesia, along with various non-motor symptoms [302]. A key pathological feature of PD is the degeneration of dopamine-producing neurons in the substantia nigra [303]. The loss of dopaminergic neurons can be modeled in a personalized whole-brain network, provided that dopamine is explicitly incorporated into the model [9]. In fact, disruptions in the dopaminergic pathways can affect communication between brain regions well beyond local activities, modifying brain dynamics as a whole [304, 305]. Abnormal bursts of activity in the beta frequency range within the basal ganglia are linked to clinical symptoms [306]. These stereotypical large-scale dynamics are associated with clinical disabilities, and interventions such as oral administration of L-Dopa and deep brain stimulation (DBS) are employed to “desynchronize” neural activity and alleviate symptoms [307, 308]. Computational modeling has explored this phenomenon both in neural network models and basal ganglia-thalamocortical circuit models [309–311]. Recent evidence combining modeling with EEG and sub-thalamic recordings has successfully simulated the spreading of brain activity between the cortex and the basal ganglia in PD [312]. In a different study, an in-silico model was fitted to 20 PD patients after DBS implantation and 15 healthy controls, demonstrating the possibility of simulating the effects of DBS stimulation from a whole-brain perspective [313]. However, the study used fMRI data (i.e., suboptimal temporal resolution), and modeled the whole cortex as a single spiking network node (i.e., suboptimal spatial resolution). In another study, virtual brain modeling approach has been used to model DBS in a proof-of-concept work, demonstrating that it is possible first to simulate pathological activity, then simulate DBS and observe: 1) restored activity at the level of the basal ganglia, and 2) large-scale changes at the cortical level [314]. Although preliminary, these studies suggest that simulating large-scale activity before and after DBS might be feasible. However, a key challenge in modeling PD is to develop computational models that include dopamine. Recently, a modular framework has been proposed to capture the effects of dopamine at the neural mass level has been proposed [99]. This new model made it possible to simulate the effect of the nigro-striatal dopaminergic tone on the brain dynamics over the large scale. When inverted, starting from individual data acquired before and after administration of L-Dopa, the model correctly predicted the dopaminergic tone (i.e., the ON or the OFF state) in each patient [9].

In stroke recovery, mechanistic models can simulate the reorganization of brain networks following an infarct. These models might help predict how different rehabilitation strategies, such as physical therapy or pharmacological interventions, could influence the re-establishment of functional connections and recovery of motor skills. As an example, it was shown that the clinical impact of different lesion patterns can be predicted utilizing brain models to quantify the distance of the dynamics from an optimal dynamical regime (i.e., near criticality) [315]. This predictive power might help design personalized rehabilitation schemes with an optimized likelihood of therapeutic success. Furthermore, the creation of virtual cohorts (that is, cohorts of virtual patients) in stroke bears promise to help the in silico testing of therapeutic strategies as well as medical equipment [11].

Besides the role in terms of patient stratification and prediction of the therapeutic outcome, computational brain models hold a special role concerning neuropsychiatric disorders [316]. Unlike other medical specialties, the nosology of neuropsychiatric ailments relies exclusively on a cluster of symptoms, as there is a lack of understanding of the underlying mechanisms. Traditional methods do not offer a way to bridge from (microscopic) mechanisms to symptoms. Computational models, however, hold promise for finding lawful associations between mechanisms and symptoms. In particular, mechanistic models allow interpreting the results and infer potential causative mechanisms [317]. These venues remain explored only superficially thus far.

Applications of VBT models have proven useful in identifying new biomarkers, even across experimental paradigms [271]. A known protocol based on TMS-EEG and the perturbational complexity index (PCI) metric, associated with consciousness levels [270], was simulated in a VBT model. A systematic exploration of the relationship between brain dynamics and PCI revealed that only one specific dynamical configuration allows complexity to be evoked by stimulation (responsiveness). Spontaneous activity was then simulated in the same VBT, and new resting-state metrics were tested, revealing that the phenomenon underlying responsiveness can also be quantified in spontaneous activity. This opens new avenues for in silico experimentation, where, with the correct mechanistic integration, seemingly independent paradigms and data can be merged into a unified theoretical framework, allowing new hypotheses to be tested with practical implications.

Computational brain models play a crucial role in advancing brain stimulation techniques. These include transcranial magnetic stimulation (TMS), deep brain stimulation (DBS), and, more recently, temporal interference (TI). These models can simulate at once the brain activities and the stimulus itself, helping to predict the effects of stimulation on different brain regions [318].

To this end, one can conceptualize the external stimulation as a perturbation that, if delivered correctly, allows the brain to reach a desired (i.e. therapeutic), state that was previously inaccessible [319, 320]. Two problems arise immediately, which can be addressed using computational models: what is the right stimulation (i.e. when and where should the brain be stimulated) and how to deliver the correct stimulation in practice [320, 321]. With respect to the latter, by understanding how electrical currents propagate through neural tissues and how they influence specific neural circuits, researchers can optimize the stimulation parameters to non-invasively deliver the necessary stimulation [322]. To this regard, TI represents a novel technique to non-invasively deliver brain stimulations [323], which bears promise as a way to induce desired behavioural changes non-invasively [324]. TI involves the application of two (or more) high-frequency electrical currents that intersect within the brain to produce a lower-frequency envelope capable of locally modulating neural activity [325]. A number of technical challenges are currently being addressed [326], such as how to generate trains of pulses of desired frequencies (and shapes) from the envelope of the amplitude-modulated signals, and how to control the shape of the stimulation focus while reducing its size (i.e. how to deliver effective, more focal stimulations) [326]. By simulating the brain’s response to these intersecting currents, computational models help researchers in understanding the spatial and temporal patterns of stimulation, and the resulting effect on the large-scale dynamics. This allows for precise targeting of specific brain regions with minimal impact on surrounding tissues. To effectively manipulate the brain network, computational models are often used within the framework of control theory [327]. With this respect, the brain is conceptualized as a dynamical system whose evolution depends on its previous dynamics, its current state, the wiring, and the characteristic of the perturbations [328]. The objective of the computational model in this context it to predict the state that the brain will access due to the stimulations, as well as the trajectory of the relaxation to the spontaneous dynamics [320, 321, 327, 328]. In neurodegenerative conditions, we recently demonstrated that perturbations in brain dynamics caused by stimulation can effectively address the non-identifiability issue in estimating the degradation of intra-hemispheric connections (within the limbic system), through increasing the probability of transitions between hidden states [226]. Through these models, researchers can determine the most desired stimulation to be delivered to a given region, and how to best engineer the pulse in order to effectively deliver it. These techniques, especially the non-invasive ones, are at their dawn, and currently a very active area of research. If successful, these approaches will enhance the precision and effectiveness of treatments of neurological and psychiatric disorders, including depression, PD, and epilepsy, leading to more personalized interventions with fewer side effects.

Currently, inference in VBTs is carried out through MCMC methods [150, 224, 228, 231], where applicable, or through SBI [225, 226] using a class of deep generative models called normalizing flows [237,329,330]. Generative models are a class of ML algorithm designed to model the underlying distribution of a dataset, enabling the generation of new, synthetic data samples similar to the original data. Techniques such as normalizing flows and variational autoencoders (VAEs; [331]) have proven effective for probabilistic inference and neural system modeling, with potential applications in diagnosis and treatment planning [7, 14, 16, 239]. Recently, Sip et al. [152] introduced a method using VAEs for nonlinear dynamical system identification at whole-brain level, which infers both the neural mass model and region- and subject-specific parameters from functional data while respecting the known network structure. Additionally, Baldy et al. [151] have applied neural ODEs [148] to spiking neurons illustrating the prediction of system dynamics and vector fields from microscopic states. These ML models are expected to enable precise, automatic inference of brain diseases from large, multimodal data in the near future.

Computational models play a critical role in the design and testing of neuromorphic devices such as brain-computer interfaces (BCIs), enabling researchers to simulate neural processes and predict how these interfaces will interact with the brain. By using these models, developers can explore different design options, optimize algorithms, and identify potential issues before implementing BCIs in real-world scenarios. For example, it has been proposed that brain models might be deployed to test causal relationships among variables of interests [332]. Furthermore, brain models can be used to optimize the design of the interfaces themselves. As an example, brain models might be used to provide a parsimonious description of the dynamics, which in turn can help define the most suitable decoders [333]. All in all, computational models might reduce the need for extensive animal or human testing, accelerate the development process, and enhance the safety and effectiveness of BCIs, paving the way for more advanced and personalized neurotechnologies. In conclusion, VBTs represent a powerful tool for predicting of therapeutic outcomes and the designing new devices that interact with the brain. Furthermore, these models offer detailed insights into the biological underpinnings of brain function and dysfunction. Unlike phenomenological models, which focus on data correlations, mechanistic models delve into the causal pathways of interventions. As our understanding of brain biology continues to deepen and brain data accumulate, mechanistic models offer a framework to generate knowledge from data and might become instrumental to more effective and personalized treatments in neurology and psychiatry [6].

## Abbreviations

VBT: virtual brain twin
TVB: the virtual brain
DTI: diffusion tractography imaging
DW-MRI: diffusion-weighted magnetic resonance imaging
CT: computed tomography
PET: positron emission tomography
BOLD: blood-oxygen-level-dependent
fMRI: functional magnetic resonance imaging
EEG: Electroen-cephalography
MEG: Magnetoencephalography
SEEG: Stereoelectroencephalography
iEEG: intracranial electroencephalography
ECoG: Electrocorticography
SC: structural connectivity
FC: functional connectivity
FCD: functional connectivity dynamic
PSD: power spectral density
NMM: Neural mass model

## Acknowledgements

This research has received funding from EU’s Horizon 2020 Framework Programme for Research and Innovation under the Specific Grant Agreements No. 101147319 (EBRAINS 2.0 Project), No. 101137289 (Virtual Brain Twin Project), and government grant managed by the Agence Nationale de la Recherche reference ANR-22-PESN-0012 (France 2030 program). The funders had no role in study design, data collection and analysis, decision to publish, or preparation of the manuscript.

## Author contributions

All authors contributed to the writing, editing, visualization, data analysis, and interpretation of the results, as well as to the overall conceptualization and development of the manuscript.

